# Effective Degrees of Freedom of the Pearson’s Correlation Coefficient under Autocorrelation

**DOI:** 10.1101/453795

**Authors:** Soroosh Afyouni, Stephen M. Smith, Thomas E. Nichols

**Author notes:** T.E.N. is the corresponding author.

## Abstract

The dependence between pairs of time series is commonly quantified by Pearson’s correlation. However, if the time series are themselves dependent (i.e. exhibit temporal autocorrelation), the effective degrees of freedom (EDF) are reduced, the standard error of the sample correlation coefficient is biased, and Fisher’s transformation fails to stabilise the variance. Since fMRI time series are notoriously autocorrelated, the issue of biased standard errors – before or after Fisher’s transformation – becomes vital in individual-level analysis of resting-state functional connectivity (rsFC) and must be addressed anytime a standardized *Z*-score is computed. We find that the severity of autocorrelation is highly dependent on spatial characteristics of brain regions, such as the size of regions of interest and the spatial location of those regions. We further show that the available EDF estimators make restrictive assumptions that are not supported by the data, resulting in biased rsFC inferences that lead to distorted topological descriptions of the connectome on the individual level. We propose a practical “xDF” method that accounts not only for distinct autocorrelation in each time series, but instantaneous and lagged cross-correlation. We find the xDF correction varies substantially over node pairs, indicating the limitations of global EDF corrections used previously. In addition to extensive synthetic and real data validations, we investigate the impact of this correction on rsFC measures in data from the Young Adult Human Connectome Project, showing that accounting for autocorrelation dramatically changes fundamental graph theoretical measures relative to no correction.

## 1. Introduction

Resting-state functional connectivity (rsFC), defined as similarity between measured brain activity between brain regions in absence of any external instructed task, has become an essential technique for understanding the human brain. Many rsFC methods make use of correlation estimated with the Pearson’s product-moment correlation coefficient 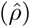, often after Fisher’s transformation (*F*) and standardised to a *Z*-score (*Z*). These *Z*-scores are used to find significant correlation or are used as a standardised measure, for example, in graph analysis where they are used to create weighted networks or are thresholded to create binary networks. However, standard results for the variance of Pearson’s correlation (before or after Fisher’s transformation) depends on independence between successive observations. Blood Oxygen Level Dependent (BOLD) time series exhibit autocorrelation which can in turn inflate the sampling variance of 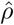. Ignoring this variance inflation – equivalently, reduction in effective degrees of freedom (EDF) – will inflate *Z*-scores and produce excess false positives when testing *H*_0_ : *ρ* = 0, and corrupt the interpretation of *Z* as a standardised effect. These biases can vary both over individuals and pairs of brain regions under consideration.

Although the impact of autocorrelation has been thoroughly investigated in task fMRI analysis (Friston et al., 2000; Woolrich et al., 2001; Lund et al., 2006), there is much less work on resting-state analyses. The first reference we are aware of directly addressing the issue in resting-state is Fox et al. (2005) which refers to “Bartlett’s Theory”, citing Watts and Jenkins (1968). Later, Van Dijk et al. (2010) uses the same approach, labeled “Bartlett’s Correction Factor”, describing it as the “integral across time of the square of the autocorrelation function”; as discussed below, this “integral” is of course a sum for discretely sampled fMRI data and, while not necessarily implied, must include both positive and negative lags of the autocorrelation function (ACF). The nominal *N* is divided by this BCF to give an EDF (see Section 2.2.1).

While these previous works use an arbitrary ACF, other authors have used an order-1 autoregressive (AR(1)) model. For example, the FSLnets toolbox (Smith et al., 2011) uses a Monte Carlo approach to estimate the variance of sample correlation coefficients (see Section 2.2.2). However, it only considers a single autocorrelation parameter over all nodes. More recently, Arbabshirani et al. (2014) made a thorough study of Pearson’s correlation variance for AR(1) time series, where, crucially, the AR(1) coefficient can vary between the pair of variables and non-null correlations were considered^1^.

Other authors have used a Wavelet representation of time series to handling autocorrelation, as Patel and Bullmore (2015) and Váša et al. (2018) use wavelet EDF-estimators initially proposed by Percival and Walden (2006) in analysis of functional connectomes obtained via wavelet transformation coefficients (Leonardi and Van De Ville, 2011).

Further, Fiecas et al. (2017) proposes an inference framework for group-level analysis of functional connectomes which accounts for the autocorrelation via the variance estimator of Roy (1989). Roy’s estimator is unique among the previous methods as it directly accounts for dependence within and between the time series, and is closely related to our method (see Section 2.2).

Alternatively, other studies have proposed pre-whitening on the resting-state BOLD time series (Christova et al., 2011; Lewis et al., 2012; Bright et al., 2017). For example, Bright et al. (2017) used pre-whitening methods, inherited from task fMRI analysis (Bullmore et al., 1996), to account for autocorrelation in restingstate analysis. Although pre-whitening is a well-established technique in task fMRI, its application in rsFC is yet to be fully investigated. Firstly, pre-whitening flattens the power spectrum which, in case of fMRI, means low frequency components are attenuated while high frequencies are amplified (Chatfield, 2016); this seems poorly suited to the resting-state analysis were the natural focus of the frequencies are on low bands, more specifically on 0.01HZ to 0.1HZ (Biswal et al., 1995). Secondly, choosing an optimal model for autocorrelation, in absence of a task paradigm, appears to be troublesome (see Bright et al. (2017)). Finally, spatial regularisation used in some neuroimaging toolboxes (e.g. FSL’s FILM) are designed for voxelwise or vertex wise analyses and would need to be adapted to Region of Interests (ROIs) data.

The concern about effect of autocorrelation on Pearson’s correlation has a long history in spatial statistics (Haining, 1991), econometrics (Orcutt and James, 1948) and climate sciences (Bretherton et al., 1999), but the fundamental work is Bartlett (1935), who first asserted that the lack of independence (between observations) is a bigger challenge than non-Gaussianity. In his 1935 paper, Bartlett proposes a variance estimator of sample correlation coefficients based on a AR(1) model, but he acknowledges that a limitation of the work is that it assumes zero cross-correlation. In later work he proposed a more general estimator which accounts for higher order AR models (Bartlett, 1946) but still fails to account for cross-correlation. Two extensions to the work has been proposed by Quenouille (1947) and Bayley and Hammersley (1946) where the former adapts the estimator to the cases where the autocorrelation functions are different for the two time series and the latter down weights the autocorrelation of long lags. Several years later Clifford et al. (1989) also proposed a reformulation of Bayley and Hammersley (1946). We have found little comparative evaluation of these methods in the literature, save Pyper and Peterman (1998) that compared False Positive Rates on low order autoregressive models of uncorrelated time series.

Importantly, save for the work of Roy (1989), all of the methods we have discussed so far have been derived under the rsFC null hypothesis (i.e. independence *between* the two series). This null encompasses both zero instantaneous and lagged cross-correlations. This is problematic for rsFC, as typically the challenge is not only to detect edges but also to measure the strength of the connectivity.

In this work we show how autocorrelation strongly influences the variance of Pearson’s correlation, breaking the variance-stabilising properties of Fisher’s transformation. We show that existing methods to adjust the variance of Pearson’s correlation for autocorrelation fail when *ρ ≠* 0, and can be severely biased if there is no or only very weak autocorrelation. To address these problems we propose a variance estimator for Pearson’s correlation that imposes no assumptions aside from stationarity, and that accounts for both auto-correlation within each time series and instantaneous and lagged cross-correlations between the time series. We call this approach “xDF”, as it comprises an effective degrees of freedom estimator that accounts for cross-correlations.

To motivate and introduce our results, as an example, we fist show how ignoring the autocorrelation may corrupt inference of correlation coefficients. Figure 1 shows the correlation of a BOLD time series in the Left Dorsal Prefrontal Cortex (PCFd) from one HCP subject to all 114 ROIs of the Yeo atlas of a *different* HCP subject (we call this inter-subject scrambling; see Section S3.6 of Supplementary Materials). Due to the random nature of resting-state BOLD time series between subjects, we expect up to 5% of the 114 correlation coefficients turn out significant (i.e. *≈* 6 regions) on average; instead, the Naive *Z*-scores (see Section 2.4) finds *≈* 35% of the regions (i.e. 40 regions) significant while xDF-corrected *Z*-scores only finds 2.6% of the regions significant (i.e. 3 regions; Figure 1.D). After application of our xDF correction, no regions survive FDR correction (2.6% significant at level 5% uncorrected). A plot of xDF-adjusted *Z*-scores against Naive *Z*-scores shows the dramatic difference in values (Figure 1.D). Observe how the connection with L-SoMotCent (blue marker) is incorrectly detected; this ROI is highly auto-and cross-correlated (blue ACF, Figure 1.E), and results in a strong correction and the *Z*-score being greatly reduced. In contrast, R-Insula (red marker) connection has essentially the same Naive and xDF-adjusted *Z*-score due to its nearly zero autocorrelation (red ACF, Figure 1.E).

**Figure 1:**
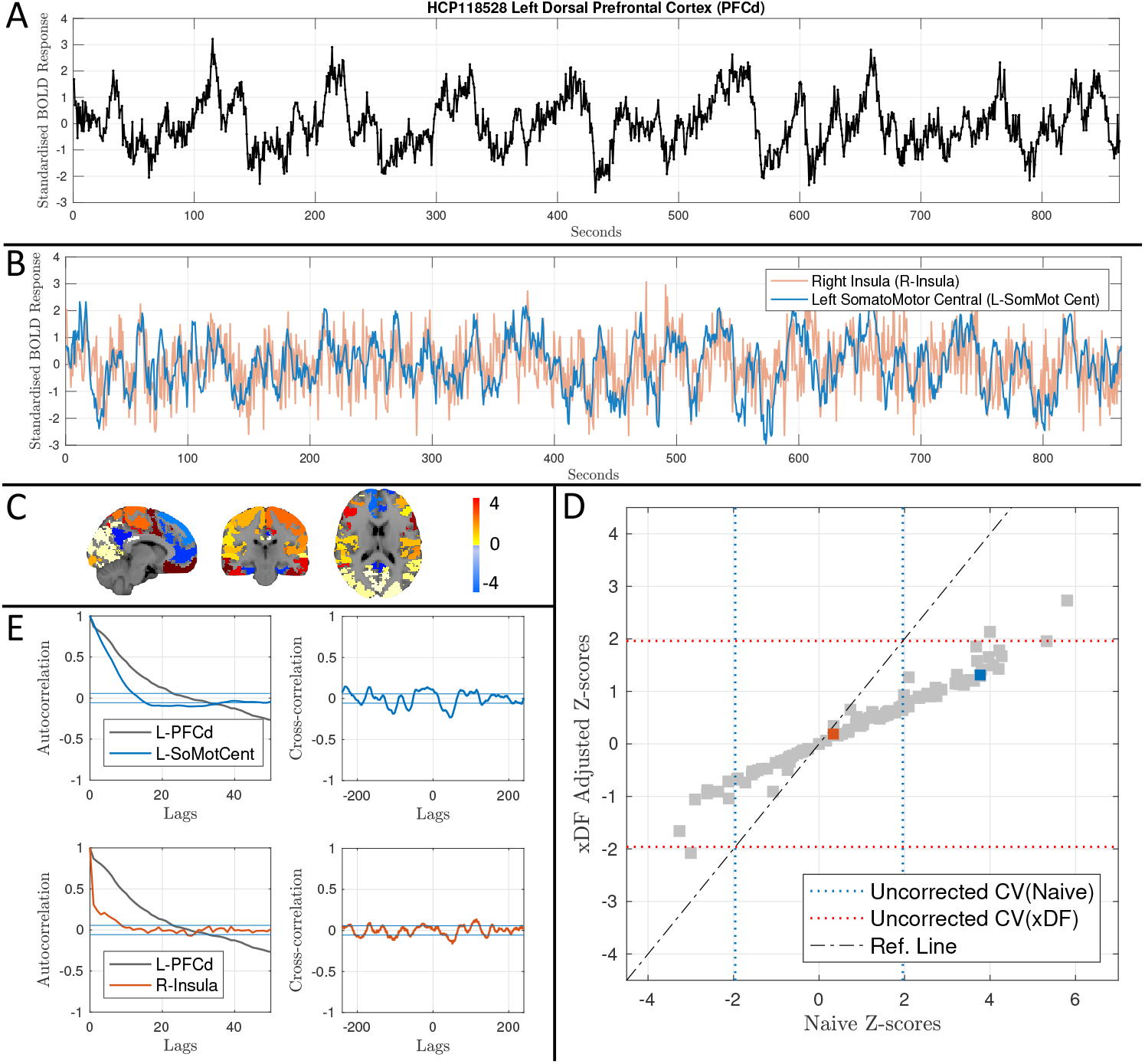
Analysis of null resting state functional connectivity to illustrate the problem of inflated correlation coefficient significance. **Panel A** shows standardised BOLD data for the Left Dorsal Prefrontal Cortex (PFCd; 421 voxles) of HCP one subject (HCP-1). **Panel B** illustrates the standardised BOLD time series of R-Insula (red; 35 voxels) and L-SomMotCent (blue; 773 voxels), illustrating the dramatically different degree of autocorrelation. **Panel C** maps the *Z*-scores of correlation between this PCFd region and time series from a *different* HCP subject (HCP-2), parcellated with the Yeo’s atlas and overlaid on an MNI standard volume. **Panel D** compares *Z*-scores accounting for autocorrelation vs. naive *Z*-scores, showing apparent significance (in this null data) with naive *Z*-scores and expected chance significance with xDF-adjusted *Z*-scores. On the horizontal axis are naive *Z*-scores that ignore autocorrelation, while on the vertical axis are *Z*-scores adjusted according to xDF. Uncorrected critical values (*±*1.96) are plotted in dashed lines. **Panel E** shows autocorrelation of each time series (left). Horizontal solid lines indicate the confidence intervals calculated as described in Section 2.3. The difference in magnitude and form of autocorrelation among the three time series is evident, with PFCd exhibiting strong, long-range autocorrelation and R-Insula showing virtually no autocorrelation. Also shown is the the cross-correlation (right panels) between HCP-1’s PCFd and HCP-2’s Left Central SomatoMotor Cortex (L-SomMotCent) (top), and HCP-1’s PCFd and HCP-2’s Right Insula (R-Insula) (bottom).

The remainder of the work is as follows. We first present a concise overview of the model and the proposed estimator. Second, we demonstrate the importance of accounting for unequal autocorrelation between each pair of variables, i.e. node-specific autocorrelation, by showing how the autocorrelation structures are spatially heterogeneous and dependent on each parcellation scheme. Third, we discuss how ignoring such effects may result in spurious significant correlations and topological features. Fourth, using Monte Carlo simulations and real data, we show how xDF outperforms all existing available variance estimators. And finally, we show how using xDF may change the interpretation of the rsFC of the human brain. The potential impact of such corrections on interpretation of rsFC is investigated for conventional thresholding method (i.e. statistical and proportional) as well as un-thresholded functional connectivity for binary and weighted networks.

## 2. Methods

### 2.1 Notation

Without loss of generality, we assume to have mean zero and unit variance time series *X* = *{x*_1_, …, *x*_*N*_ *}* and *Y* = *{y*_1_, …, *y*_*N*_ *}*, each of length *N*, with 𝕍(*X*) = **Σ**_*X*_ and 𝕍(*Y*) = **Σ**_*Y*_ ; we write the cross-correlation matrix between *X* and *Y* as 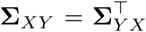. We assume stationarity, and thus have Toeplitz **Σ**_*X*_, **Σ**_*Y*_ and **Σ**_*XY*_, and denote autocorrelation of *X*

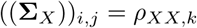

Let *i* and *j* be row and column of the covariance matrix, then *k* = *i - j*, and likewise for *Y*. The cross-correlations between *X*’s time point *i* and *Y* ‘s time point *j* is

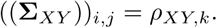

The key rsFC parameter is *ρ*_*XY*,0_, the cross-correlation at lag 0, which we refer to as simply *ρ* going forward. Note that the cross-correlation matrix is not symmetric, and so *ρ*_*XY,k*_ and *ρ*_*XY,-k*_ are distinct.

### 2.2 Variance of Sample Correlation Coefficients

For the sample correlation coefficient of mean zero data, 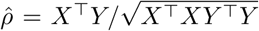, we can derive a general expression for its variance (Appendix B):

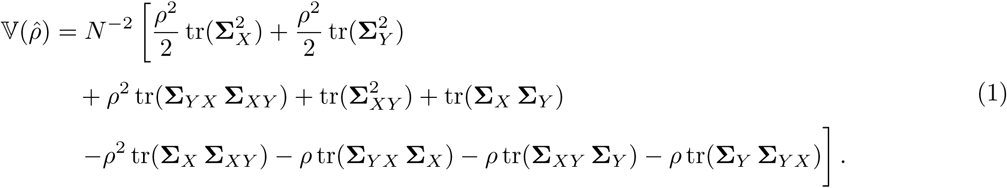

For a stationary covariance (see Appendix C), we can rewrite Eq. 1 as

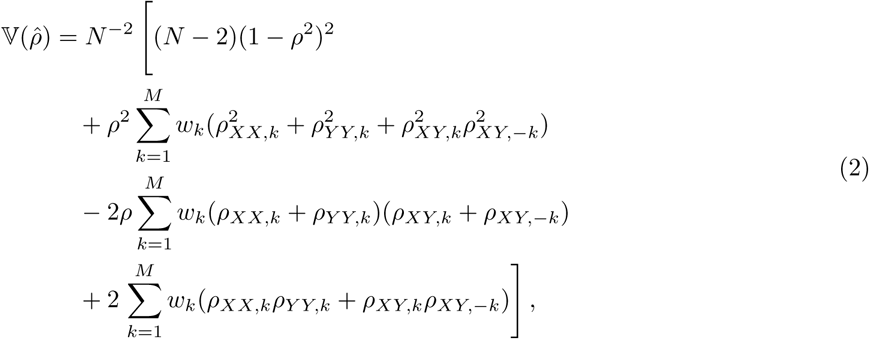

where *w*_*k*_ = *N −* 2 *− k*. While Eq. 2 takes the same form as the estimator of Roy (1989), we obtain our result from finite sample as opposed to asymptotic arguments (see Appendix B).

It is also useful to discuss two special cases of the Eq. 1. First, suppose two time series *X* & *Y* are both white but correlated such that **Σ**_*X*_ = **Σ**_*Y*_ = **I**, and **Σ**_*XY*_ = **I***ρ*. Eq. 1 then reduces to

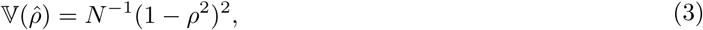

the well-known result for the variance of the sample correlation coefficients between two white noise time series (see Lehmann (1999b), §5.4).

Second, suppose *X* and *Y* are autocorrelated but are uncorrelated of each other, with non-trivial **Σ**_*X*_ and **Σ**_*Y*_ but **Σ**_*XY*_ = **0**. Then, Eq. 1 reduces to

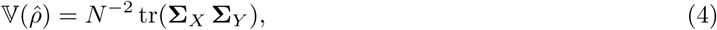

a result on the variance inflation of *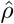* proposed by Clifford et al. (1989) and also discussed as the variance of the inner product of two random vectors in Brown and Rutemiller (1977). Written in summation form (see Appendix C) this expression is

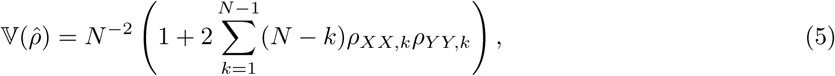

which is a result proposed much earlier by Bayley and Hammersley (1946) and which has found use in neuroimaging (Nicosia et al., 2013; Valencia et al., 2009). A closely related form (Dutilleul et al., 1993) that adjusts for mean centering has also been used in neuroimaging (Nevado et al., 2012; Pannunzi et al., 2018), though for typical time series lengths (i.e. *N ≫* 20) there should be little difference from the original result.

#### 2.2.1 Effective Degrees of Freedom for the Correlation Coefficient

One way of dealing with autocorrelation is to modify a variance result that assumes no autocorrelation, replacing *N* with a deflated EDF 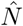. This can be done in terms of 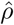 (e.g. Eq. 3) or after Fisher’s transformation; here we consider EDFs for 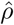 and return Fisher’s transformation in Section 2.4.

Different corrections have been proposed to estimate 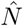. One of the earliest results is due to Bartlett (1935), who proposed an EDF for uncorrelated (*ρ* = 0) AR(1) time series:

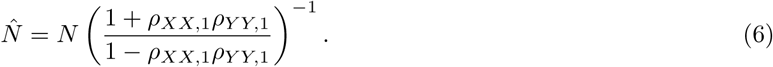

We refer to this EDF estimator as B35.

Building on work of Bartlett (1946), Quenouille (1947) proposed a more general EDF that allowed for any form of autocorrelation,

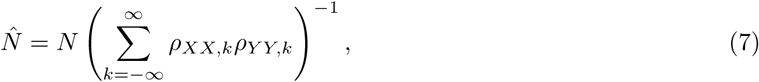

though still assuming *ρ* = 0. We refer to this EDF estimator as Q47.

In neuroimaging, a *global* form of Eq. 7 has been used, where a single ACF *ρ*_*GG,k*_ is computed averaged across voxels or ROIs for each subject, or even over subjects (Fox et al., 2005; Van Dijk et al., 2010); it takes the form

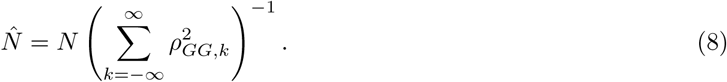

We refer to this EDF as G-Q47.

Finally, the variance result due originally Bayley and Hammersley (1946) and Clifford et al. (1989) (Eqs. 4 & 5), gives EDF

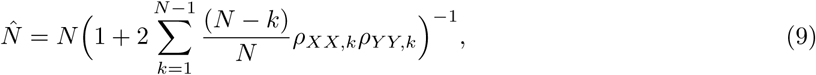

still under an independence assumption *ρ* = 0. We refer to this EDF as BH.

Whether defined with infinite or finite sums, some sort of truncation or ACF regularisation is required to use these results in practice, which we consider in Section 2.3.

#### 2.2.2 Monte Carlo Parametric Simulations

The one other approach we evaluate is Monte Carlo parametric simulation (MCPS) (Ripley, 2009). In this approach the variance of the sample correlation is estimated from surrogate data, simulated to match the original data in some way. If a common autocorrelation model and parameters are assumed over variables and subjects, this can be a computationally efficient approach. For example, the FSLnets^2^ toolbox for analysis of the functional connectivity assumes an AR(1) model with the AR coefficient chosen globally for all subjects and node pairs. While MCPS avoids any approximations for a given model, it can only be as accurate as the assumed model.

We evaluate the method used by FSLnets, which chooses the number of realisations set equal to the number of nodes. We refer to this as AR1MCPS.

### 2.3 Regularising Autocorrelation Estimates

All of the advanced correction methods described depend on the true ACFs *ρ*_*XX,k*_ and *ρ*_*Y Y,k*_, and some on the cross-correlations *ρ*_*XY,k*_. We expect true ACFs and cross-correlations to diminish to zero with increasing lags, but sampling variability means that non-zero ACF estimates will occur even when the true values are zero. Thus all ACF-based methods use a strategy to regularise the ACF, by zeroing or reducing ACF estimates at large lags.

Several arbitrary rules have been suggested for truncating ACF’s, zeroing the ACF above a certain lag. For example, Anderson (1983) suggests that the estimators should only consider the first 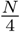 lags or Pyper and Peterman (1998) has found that truncating at 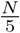 lags is optimal. Since the latter study provides a thorough empirical evaluation, we choose 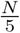 as the cut-off lag for methods Q47 and B46.

For the xDF method we considered a range of regularisation approaches. Smoothly scaling ACF estimates to zero with increasing lag is known as tapering. Chatfield (2016) suggests tapering methods using Tukey or Parzen windows. For example, for Tukey tapering, the raw ACF estimate is scaled by 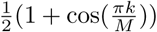 for *k <*= *M* and zeroed for *k > M*. Similar to truncating, finding the optimal *M* appears to be cumbersome; Chatfield (2016) suggests an *M* of 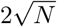 while Woolrich et al. (2001) propose the more stringent 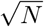; for a detailed comparison of tapering methods in fMRI see Woolrich et al. (2001).

For computation of the xDF correction we consider fixed truncation and Tukey tapering, as well as an adaptive truncation method. For our adaptive method, we zero the ACF at lags *k ≥ M*, where *M* is the smallest lag where the null hypothesis is not rejected at uncorrected level *α* = 5%, based on approximate normality of the ACF and sampling variance of 1*/N*. We base the truncation of the cross-correlation *ρ*_*XY,k*_ on the ACFs of *X* and *Y*, choosing the larger *M* found with either time series. Unless stated otherwise, the adaptive truncation method is used with xDF. For summary of regularisation methods, please see Table 2.

### 2.4 Fisher’s Transformation

It is typical to apply Fisher’s transformation to correlation estimates, 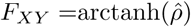, which has approximate variance

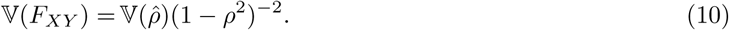

Fisher’s transformation is derived to cancel the effect of *ρ* on 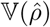 in the *absence of autocorrelation*; recall 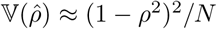 (Eq. 3) for the no-autocorrelation case. (Fisher derived a more precise variance in this setting, 𝕍(*F*_*XY*_) = (*N −* 3)^*-*1^, but this is yet still an approximation (Fouladi and Steiger, 2008).) In the presence of autocorrelation, the variance of *F*_*XY*_ remains dependent on *ρ* (more generally Σ_*XY*_, as well as Σ_*X*_ and Σ_*Y*_) and *F*_*XY*_ cannot be regarded as a variance-stabilised quantity. Only with an accurate estimate of 𝕍(*F*_*XY*_), that considers auto-and cross-correlation, can the final *Z*-score be considered “stabilised”.

We focus the remainder of our evaluations on *Z*-scores of the form

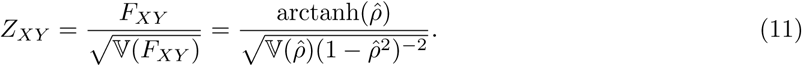

Each particular correction method used determines 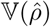. For xDF we use Eq. 1, while for all other methods we use the nominal variance with an EDF, i.e. 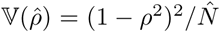; Naive has 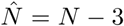, and each other EDF method uses their respective estimate 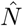, as described in Section 2.2.1 and Table 1.

**Table 1:** Summary of the EDF 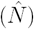 estimators for sample correlation coefficients. For simplicity, we refer each method by initials of the authors.

| Method | Equation | Reference | Neuroimaging Applications |
| --- | --- | --- | --- |
| Naive | $\hat{N} = N - 3$ | Fisher (1915) | Bassett et al. (2011)<br>& Gonzalez-Castillo et al. (2014)<br>& Cao et al. (2014) |
| B35 | Eq. 6 | Bartlett (1935) | Not found |
| BH | Eq. 4 | Bayley and Hammersley (1946) &<br>Clifford et al. (1989) | Nicosia et al. & Valencia et al. (2009)<br>Nevado et al. (2012) & Pannunzi et al. (2018)<br>Simas et al. (2015) |
| Q47 | Eq. 8 | Quenouille (1947) & Bartlett (1946) | Fox et al. (2005) & Zhang et al. (2008) |
| xDF | Eq. 1 & 2 | Roy (1989) & the current work | Fiecas et al. (2017) |
| AR1MCPS | Sec. 2.2.2 | Smith et al. (2011) & Ripley (2009) | Nomi and Uddin (2015) Smith et al. (2015) |

**Table 2:** Summary of the regularisation methods for autocorrelation function of time series *X*.

| Method | Cut off lag | Window Type | Reference |
| --- | --- | --- | --- |
| Truncation | $M = N/4$ &<br>$M = N/5$ | N/A | Anderson (1983) &<br>Pyper and Peterman (1998) |
| Adaptive Truncation | $H_0 : \rho_{XX,k} = 0$ | N/A | N/A |
| Tapering | $M = \sqrt{T}$ &<br>$M = 2\sqrt{T}$ | Parzen & Single Tukey | Chatfield (2016) &<br>Woolrich et al. (2001) |

### 2.5 Simulations and Real Data Analysis

The xDF is validated and compared with other existing estimators via series of Monte Carlo simulations and real data experiments. We simulate time series with various autocorrelation structures (see Section S3.1), under both uncorrelated and correlated conditions, using ACF parameters estimated from one HCP subject (see Section 3.4). We generate null realisations with real data by randomly exchanging the nodes between subjects (see Section S3.6). From both of these sources of null data we evaluate the distribution of *Z*-scores and false positive rates.

To investigate sensitivity and specificity, we simulate correlation matrices, transformed to *Z*-scores with each method, with 15% of edges considered as signal (i.e. assigned with *ρ >* 0). Briefly, sensitivity refers to the proportion of true positives (i.e. edges which were assigned with a non-zero correlation and also rejected null hypothesis, *H*_0_ : *ρ* = 0) over all positives (i.e. all edges which have rejected *H*_0_). Specificity is defined as proportion of true negatives (i.e. edges which were assigned with zero correlation and also failed to reject *H*_0_) over all negatives (i.e. all non-detected edges). Accuracy is defined as the summation of two measure described (see Section S3.5).

We consider graph metrics computed on real data, based on *Z*-scores from each method. We use one session of resting state data from each of the 100 unrelated HCP subjects. This data was pre-processed (Section S4) and we created *P × P* resting-state functional connectivity matrices (*Z*-scores), where *P* is number of ROIs, depending on the choice of parcellation scheme; we use the Yeo, Power and Gordon atlas in their volumetric form and ICA200 and Multimodal Parcellation (MMP) in surface mode (see Section S4.1). The rsFC matrices were then thresholded using two conventional thresholding methods; statistical thresholding (where FDR corrected *α* = 5% is used to test the significance of each edge; see Section S4.3) and proportional thresholding (where matrices are thresholded on population cost-efficient density so all matrices have identical density; see Section S4.4). Finally, the effect of the autocorrelation corrections on two centrality measures (weighted degree and betweenness) and two efficiency measures (local and global) are investigated (see Section S4.5.3).

In all our evaluations we estimate the ACFs from the time series, incorporating this important source of uncertainty (Algorithm S2 in Supplementary Materials). An exception are “Oracle” simulations in which the true ACF parameter values are used when estimating the variance (see Algorithm S1 in Supplementary Materials).

### 2.6 Autocorrelation Index

To summarise the strength of autocorrelation in time series *X*, we use

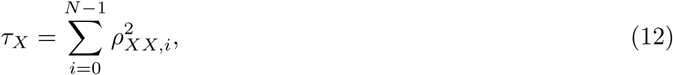

which we call the autocorrelation index (ACI).

## 3. Results

### 3.1 Autocorrelation and Parcellation Schemes

We find that the degree of autocorrelation of resting state data is highly heterogeneous over the brain. Figure 2.A shows a maps of voxel-wise and ROI ACI, averaged across subjects (15 for voxelwise, 100 for ROIs), showing that ACI vary widely. We found that using parcellation schemes not only fails to reduce the spatial heterogeneity, but instead magnifies the autocorrelation effects: Figure 2.B shows autocorrelation indices for three ROIs of Yeo’s atlas, for each voxel in an ROI and as ROI averages: Left posterior cingulate (LH-PCC; 1091 voxels), Left somatosensory motor (LH-SomMot; 4103 voxels) and Left dorsal prefrontal cortex (LHPFC; 19 voxels). We find a dramatic increase in autocorrelation with averaging within ROIs (see Appendix E for the likely origin of this effect).

**Figure 2:**
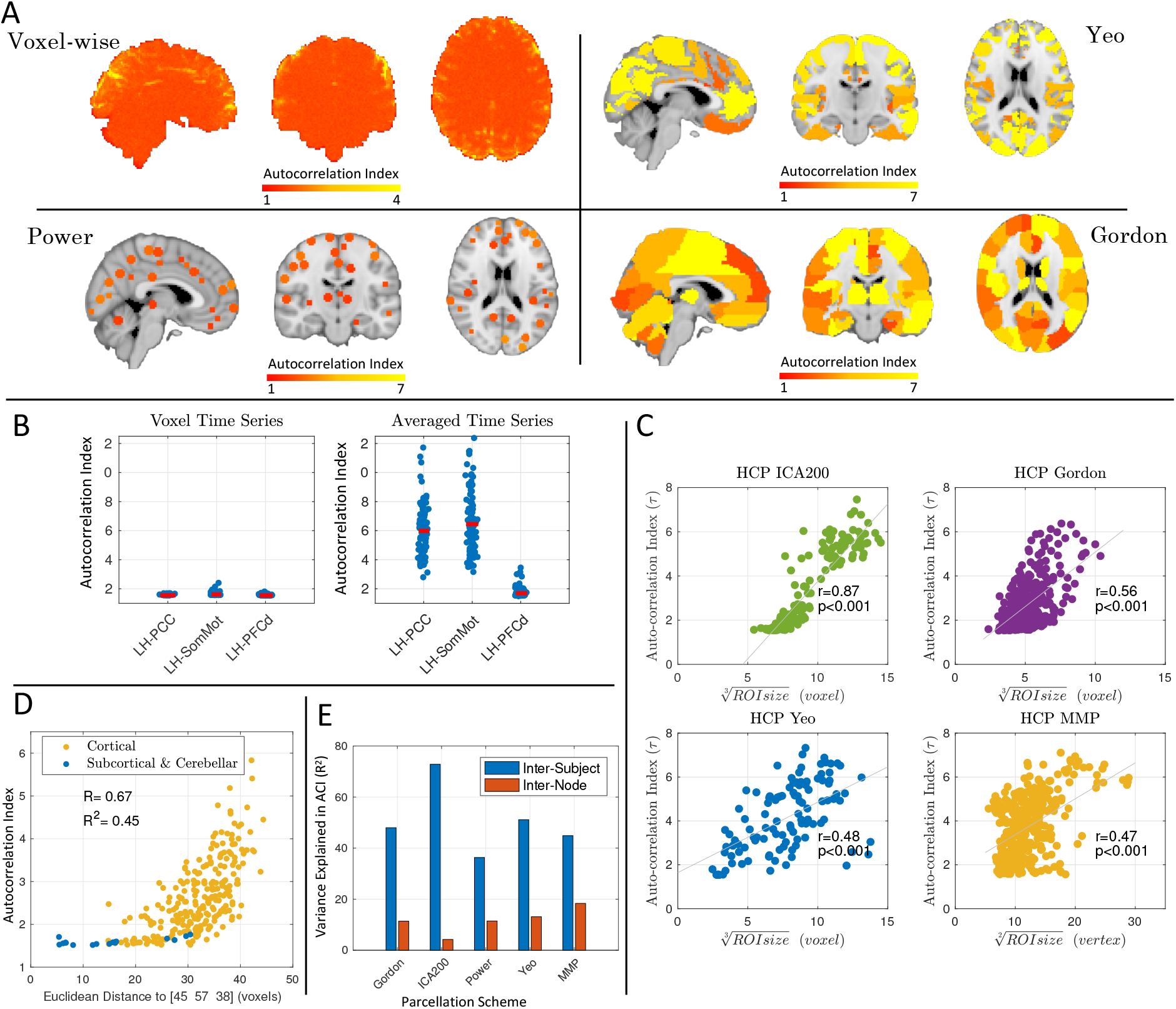
Variation in strength of autocorrelation over space within an atlas, and between atlases. **Panel A** maps the autocorrelation index (ACI) voxelwise and for 3 different atlases, averaged over subjects (15 for voxel-wise, 100 for ROIs); variation is particularly evident for Yeo and Gordon; Power atlas is more homogeneous (but see Panel D). **Panel B** shows the impact of averaging within ROIs on autocorrelation. Left, shows ACI of individual voxels (blue dots) of a single subject across three regions of interests (ROIs) from the Yeo atlas. Right panel illustrates the ACI of ROI-averaged time series (blue dots) for 100 subjects, showing dramatic increase in ACI; red lines indicates the median. ROIs are Left Posterior Cingulate (LHPCC), Left Somatosensory Motor (LH-SomMot) and Left Dorsal Prefrontal Cortex (LH-PFC). **Panel C** plots the ACI, averaged over subjects of the HCP 100 unrelated-subjects package, vs. region size for ACI time series and three atlases, where ICA and one atlas (MMP) are surface-based. There is a strong relationship between ACI and ROI size. The “ROI size” for ICA is defined as number of voxels in each component above an arbitrary threshold of 50. For MMP, the ROI size is defined as number of vertices comprising an ROI. **Panel D** considers the Power atlas, which has identically sized spherical ROIs, plotting ACI vs. distance to a voxel in the thalamus. Cortical ROIs have systematically larger ACI than subcortical ROIs. **Panel E** shows variance explained by inter-subject and inter-node ACI profiles for the Gordon, ICA200, Power and Yeo atlases; the large variance explained by inter-subject mean indicates substantial consistency in ACI over subjects.

More generally, we find that the size of ROIs predicts autocorrelation of the ROI in both volumetric and surface-based parcellation schemes (Figure 2.C). Further, not only the size of the ROI, but the location of the ROI influences autocorrelation. Using the Power atlas, where all ROIs have identical volume (81 2mm^3^ voxels), the autocorrelation in subcortical structures is weaker than in cortical structures, as summarized by plotting autocorrelation index vs. distance to Thalamus (Figure 2.D). While differences in BOLD characteristics between subcortical and cortical voxels could contribute to the autocorrelation structure, it is more likely the higher noise levels (far from the surface coils, and more susceptible to problems of acceleration-reconstruction, in this HCP data) explain the lower autocorrelation index in subcortical regions.

Using an ANOVA with either node or subject as the explanatory variable, we quantify the heterogeneity of autocorrelation index as the variance explained by variable (Figure 2.E). For time series extracted using Power atlas, 36% of inter-subject ACI variance is explained, while for ICA200 time series up to 73% of inter-subject is explained, showing that the severity of autocorrelation is very subject-dependent regardless of atlas used. The ACI variance explained by node is smaller, but above 12% for all four parcellation schemes, suggesting the importance of node-specific autocorrelation adjustment.

### 3.2 Real Data and Monte Carlo Evaluations

We use inter-subject scrambling of 100 HCP subjects, parcellated with Yeo atlas, to create null realisations with realistic autocorrelation structure. Using these realisations we compare different EDF methods in terms of FPR and distribution of Fisher’s *Z*-scores, both visually via QQ plots and by Kolmogorov–Smirnov (KS) statistics of observed *Z*-scores vs. a standard normal distribution. Results on node-specific autocorrelation corrections in Figure 3A show that Naive and B35 have greatly inflated FPR, while BH and its approximation, Q47, successfully preserve the FPR level at the 5% level while distribution of the both methods closely follow normal distribution (i.e. -log10(KS) = 2.54). Similar results for other FPR levels (%1 and 10%) are also illustrated in Figure S4.

**Figure 3:**
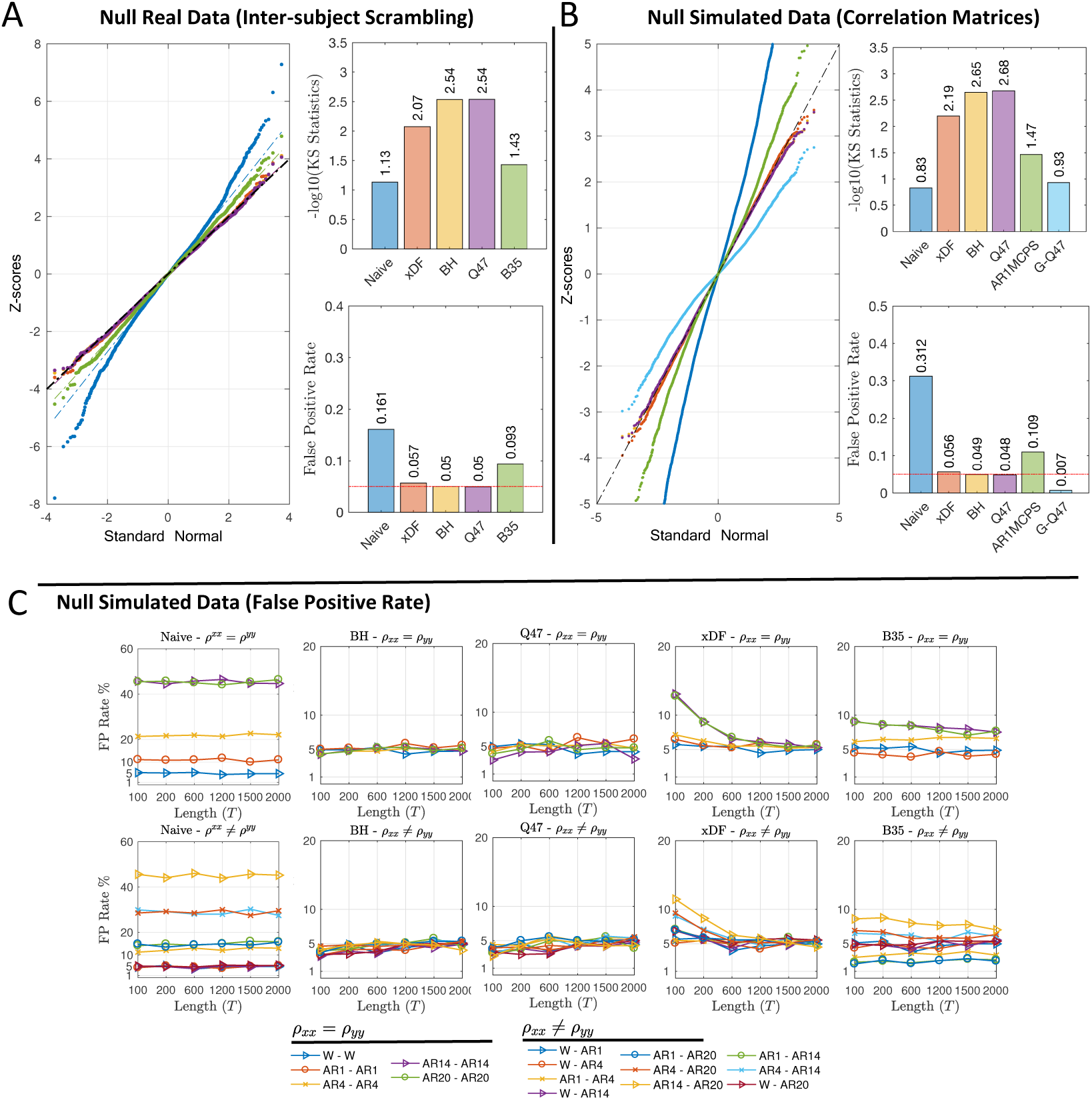
Evaluation of false positive rate control for testing *ρ* = 0 with different autocorrelation correction methods. **Panel A** shows results using real data and inter-subject scrambling of HCP 100 unrelated subjects with the Yeo atlas ROIs, comprising 235,500 distinct *Z*-scores (see Figure S6 for same results with other atlases). Left shows the QQ plot of *Z*-scores of each method, top right shows the *-* log10 KS statistics (larger is better, more similar to Gaussian), and bottom right the FPR, all of which show that Naive and B35 have very poor performance. **Panel B** depicts a similar evaluation with simulated data, where a single ACF is used to simulate all time series with identical autocorrelation (see Section S3.5), again under the null; we additionally consider two “global” correction methods that assume common ACF between the nodes, G-Q47 and AR1MCPS. Here the Naive and the two global methods have poor false positive control. **Panel C** shows the FPR at the nominal 5% *α* level across five methods (columns) for identical (top row) and different (bottom row) ACFs, over a range of time series lengths. Naive (note different y-axis limits) and B35 have poor FPR control, while BH, Q47 and xDF all have good performance for long time series, with xDF having some inflation for the most severe autocorrelation structures with short time series. The setting of each simulation is coded by plotting symbol and colour, as shown at the bottom of the figure.

Since the majority of methods used are not node-specific but instead consider the autocorrelation as a global effect (Fox et al., 2005; Zhang et al., 2008; Hale et al., 2016), we evaluate them under their homogeneity assumption using simulated correlation matrices comprised of uncorrelated time series with strong autocorrelation measured from one particular HCP subject (see Section S3.5 for details). Figure 3.B shows comparison of node-specific methods (B35 excepted, due to its poor FPR control) to global methods AR1MCPS and G-Q47. Both global correction methods fail to achieve the desired FPR level and the KS statistics of the two methods are also amongst the lowest; this poor performance is likely due to the simple AR correlation model used by each of these methods. On the other hand, the node-specific methods (xDF, BH and Q47) remarkably improves the FPR and KS statistics. However, for uncorrelated time series, BH and GQ47 outperform the xDF. Similar results for other FPR levels (%1 and 10%) are illustrated in Figure S5.

We also repeated the FPR and KS analysis of correlation matrices for another set of simulated correlation matrices, this time with autocorrelation structures drawn from a different subject than above. Results suggest similar FPR and KS statistics except for AR1MCPS which almost meet the FPR-level. This clearly suggests that the performance of the global measures, especially AR1MCPS, are subject-dependent.

We further complement the validation methods with FPR and ROC analysis. Using the same simulation techniques, discussed in section S3.1, we compare the FPR levels for pair-wise uncorrelated time series. Figure 3.C illustrates the FPR of each method for level *α* = 5%. Figure 3.C suggests that the Naive correction (first column) can only maintain the desired FPR level when at least one of the time series are white, otherwise, the FPR level can approach 50%, in cases where the both time series are highly autocorrelated. Second and third columns of Figure 3.C shows the FPR levels after the degrees of freedom are corrected via BH and Q47 methods where results suggest a remarkable improvement. Despite both methods having a conservative FPR level on short time series, both successfully maintain the FPR level on larger *N*. The FPR results for xDF suggest that for short time series, the method fails to contain the FPR level, especially on highly autocorrelated time series, however as *N* grows, the FPR levels approaches the nominal level *α* until *N* = 2000 where the xDF has the closest FPR level. We finally, evaluate the FPR of B35 where, for majority of the autocorrelation structures, the method has failed to control the FPR level regardless of the sample size. For example, for time series with AR1-AR14 structure, the FPR level is as conservative as 2% while for AR14-AR20 the level exceeds 7%.

Results presented in Figure 3 are for highly autocorrelated, yet uncorrelated, time series (*ρ* = 0). However, in rsFC, it is the highly correlated time series that are of interest. This motivates us to investigate the accuracy of standard errors for 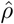 for highly autocorrelated time series, with non-zero cross-correlation, simulated following the model described in Section S3.1.

The bias for estimating Pearson’s correlation standard deviation 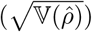 by the Naive method is severe and varies with *ρ* when the ACF’s are unequal (Figure S2). For the other methods, Figure 4.A, 4.B & 4.C show percent bias for existing methods, B35, BH and Q47, respectively. While BH and Q47 corrections give mostly unbiased standard errors in case of independence (*ρ* = 0) there is substantial bias as correlation *ρ* grows, for both short and long time series length. For example, for *ρ* = 0.5, BH and Q47 corrections overestimates 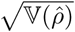 by more than 30%. The bias for the B35 standard errors show a similar pattern but with particular sensitivity to the autocorrelation structure.

**Figure 4:**
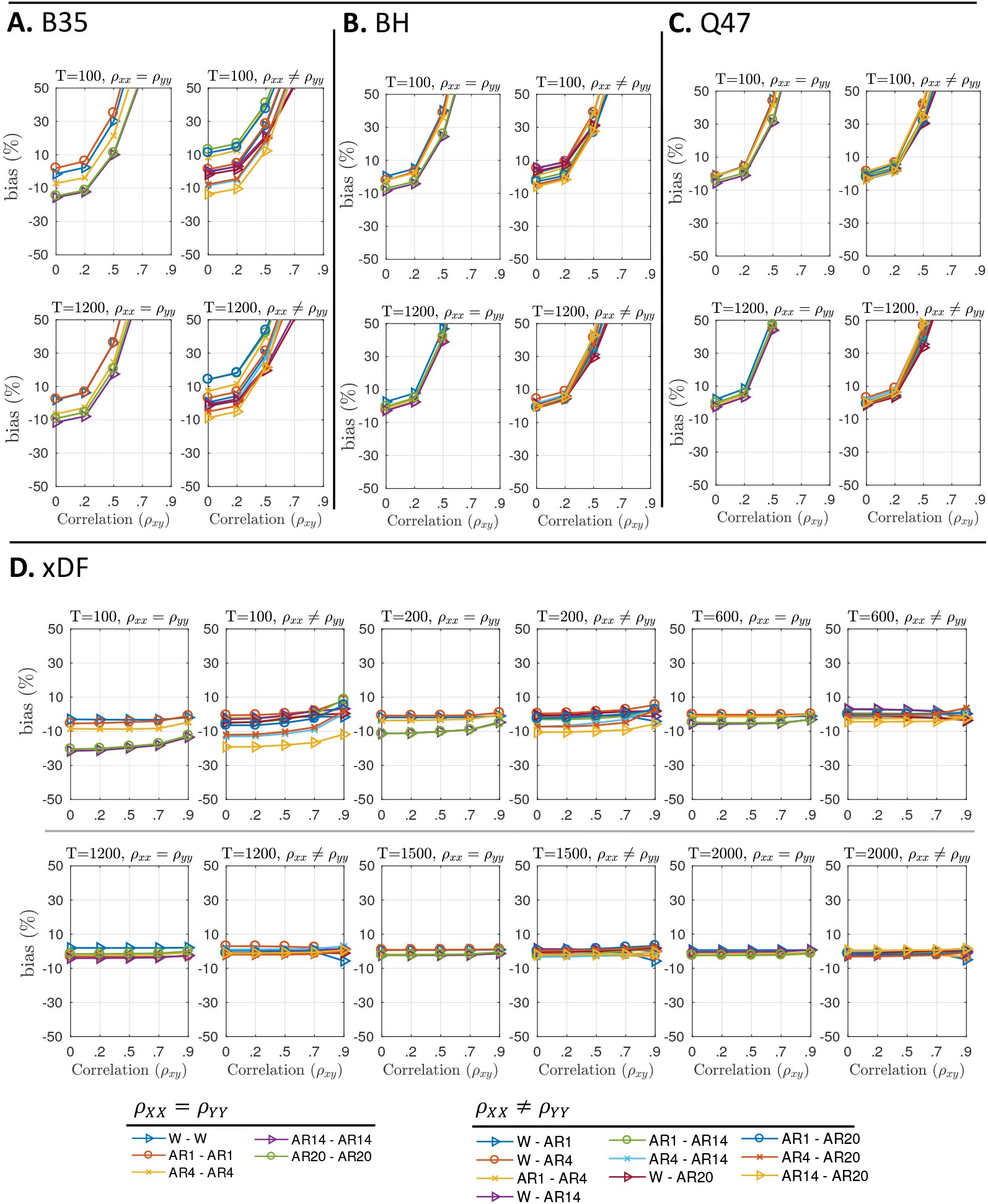
Percentage bias of estimated standard deviation of 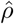 for different autocorrelation correction methods. **Panel A** plots the bias of the B35 method for *T* = 100 (top) and *T* = 1200 (bottom) for equal (left) and unequal (right) ACF’s. **Panel B** plots the same for BH, and **Panel C** for Q47. **Panel D** plots the same information for a wider range of time series lengths *T*. These results show the dramatic standard error bias in BH35, BH and Q47 with increasing *ρ*. All results here are for our adaptive truncation method; see Figures S8 and S9 for percent bias of different tapering methods. The setting of each simulation is coded by plotting symbol and colour, as shown at the bottom of the figure. Details of simulations and bias computation are found in Supplementary Materials; see Algorithm S2 and Eq. S10. We exclude the results for biases of Naive standard error as they often exceed up to %60 for autocorrelated time series; see Figure S2

A notable finding from these *ρ ≠* 0 results is for white time series (“W-W” for *ρ*_*XX*_ = *ρ*_*Y Y*_, blue triangles). For B35, BH and Q47 methods, this ‘easy’ case of no autocorrelation gives just as bad performance as severe autocorrelation. We identified the source of this problem as a confounding of the product of sample autocorrelation functions with sample cross-correlation; see Appendix D for details.

For xDF (Figure 4.D), the performance is dramatically better, with less 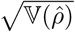 bias over all and no notable dependence on *ρ*. The worst performance is for short time series and high-order AR autocorrelation, but for *N ≥* 200 bias is mostly within *±*5% and improves with *N*.

Results for Oracle simulation (Figure S3.A) also confirm the confounding of autocorrelation with cross-correlations, as Q47 and BH are both biased even when the true parameters are used, while xDF shows negligible biases across different autocorrelation structures and sample sizes.

We also use the simulations to evaluate *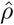* xDF’s standard error bias across different tapering methods (see section 2.3). Figure S8 suggests that despite similarities between the tapering methods on low-and mid-range correlations, they differ on higher correlation coefficients where unregularised, Tukey tapered 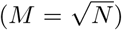 and truncation (*M* = *N/*5 lags) autocorrelation functions overestimate the variances while the shrinking and Tukey of 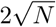 lags maintain the lowest biases. Although the two methods, Tukey taper with cut-off at 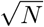 lags and adaptive truncation, appear to have very similar biases we notice that adaptive truncation has less bias for short time series. Moreover, adaptive truncation is immune from arbitrary choice of lag cut-off. We therefore use adaptive truncation as the ACF regularisation method for remainder of this work.

The FPR analysis presented in Figure 3.C concern only the null case of uncorrelated time series. To summarise performance in the presence of correlation *ρ >* 0 we evaluate the sensitivity and specificity of each method, via ROC analysis, on simulated correlation matrices, discussed in section S3.5, where the time series are highly dependent and autocorrelated. Figure 5 illustrates the sensitivity, specificity and accuracy measures for each of the methods across three different sample sizes. For correlation matrices comprised of both short and long sample sizes, the xDF outperforms other methods in terms of accuracy. While other measures have higher sensitivity than xDF, they suffer from worse specificity. AUC analyses showed virtually no difference between the methods for FPR *<* 10% (Fig. S7).

**Figure 5:**
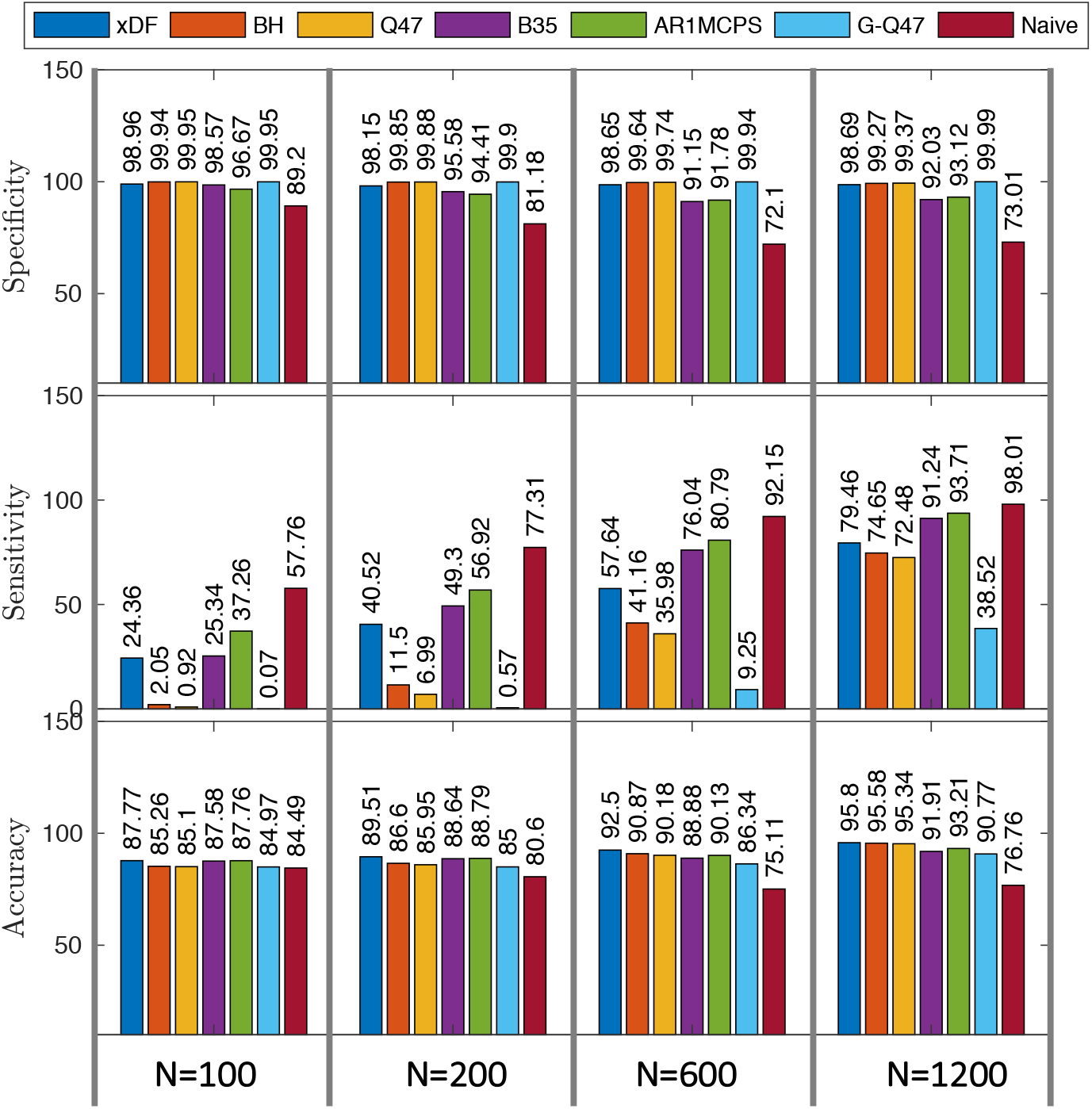
Performance of testing *ρ* = 0 at level *α* = 0.05 on 5000 simulated correlation matrices (114 × 114, matching Yeo atlas) with 15% non-null edges (see Section S3.5). From top to bottom, specificity, sensitivity and accuracy (sum of detections at non-null edges and non-detections at null edges) are shown. Specificity (i.e. FPR control) is good for xDF, BH, Q47 and G-Q47, and sensitivity increases with time series length; accuracy is best for xDF, closely followed by BH and Q47.

### 3.3 Effect of Autocorrelation Correction on Functional Connectivity

Figure 6.A suggests that, as expected, the *Z*-score of the functional connections either remained unchanged or has been reduced due to unbiased estimation of variance using xDF correction. For example, the functional connection between node 37 and 94 (i.e. both series are almost white; see Figure 1.E) has experienced almost no changes; *Z*_xDF_(37, 94)=3.61 and *Z*_Naive_(37, 94)=3.65, while functional connection between nodes 23 and 13 (i.e. both series are highly autocorrelated; see Figure 1.E) was reduced for 200%; *Z*_xDF_(13, 25)=2.08, *Z*_Naive_(13, 25)=6.02.

**Figure 6:**
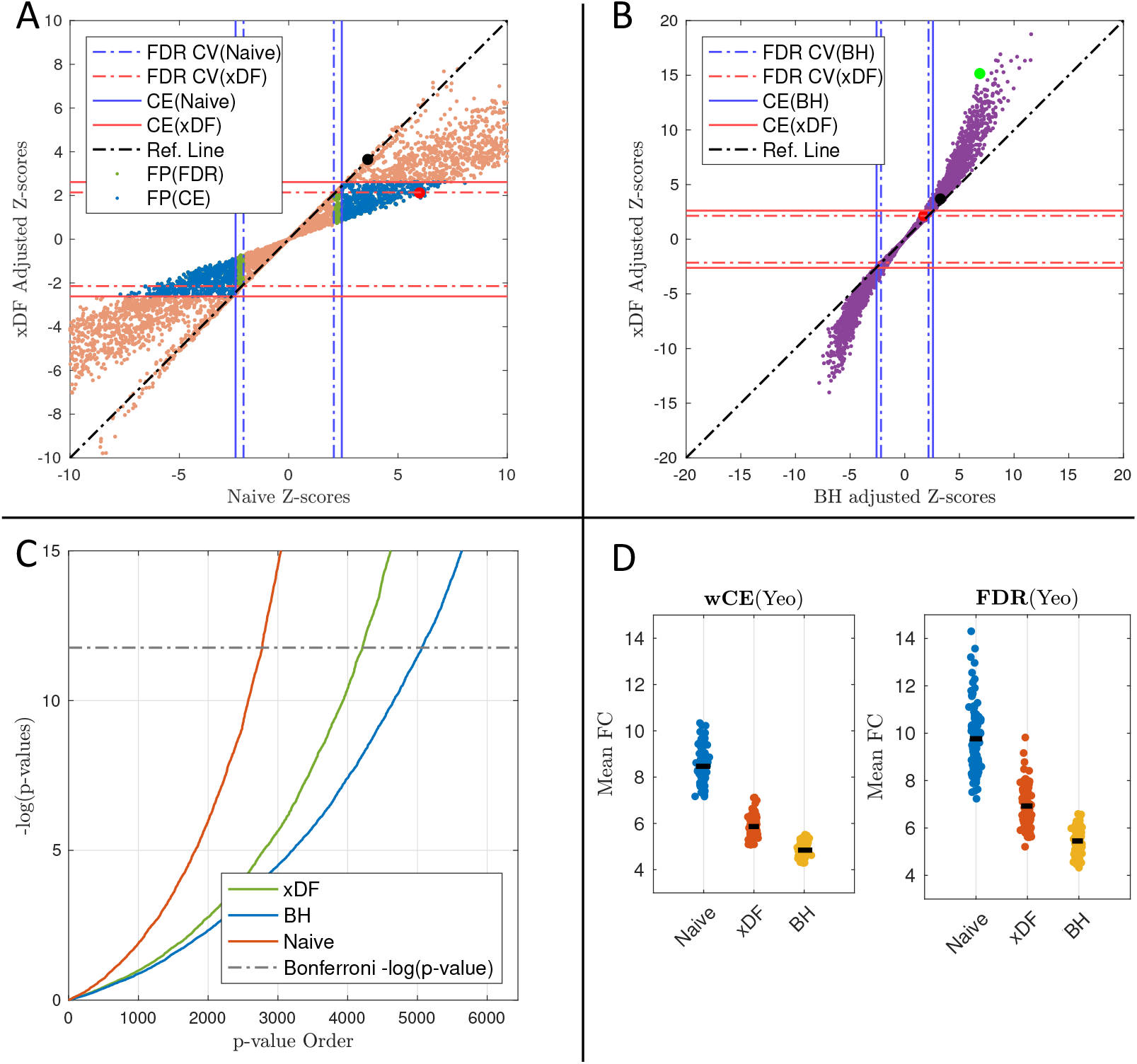
Impact of Naive, xDF and BH corrections on rsFC in one HCP subject parcellated with the Yeo atlas. **Panel A** plots rsFC *Z*-scores of xDF-corrected connectivity vs. Naive, showing that the significance of edges with Naive computation of *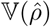* is almost always inflated, but to varying degrees. Solid lines are the critical values corresponding to the cost-efficient (CE) density. Dashed lines illustrates the critical values of FDR-corrected q-values. Taking xDF as reference, edges that are incorrectly detected with Naive are coloured green (FDR but not CE) and blue (CE). The black point marks edge (37,94) and red point (13,25), discussed in body text. **Panel B** plots rsFC *Z*-scores of xDF-corrected connectivity vs. BH, same conventions as Panel A, showing deflated significance of *Z*-scores computed with the BH method. The green point marks edge (103,104). **Panel C** pp-plot of p-values of *Z*-scores from xDF (green), BH (blue) and Naive (red) corrections. Dashed line is %5 Bonferroni threshold for 6,441 edges. **Panel D** shows the differences in mean functional connectivity (mFC) of each correction method across statistical (FDR) and proportional (CE) thresholding. See Figure S10 for a similar plot for a different HCP Subject.

Naturally, such drastic changes in *Z*-scores are also reflected in p-values of statistical inferences for each connection. Figure 6.C illustrate these changes (i.e. orange dots) between FDR-corrected p-values (i.e. q-values) of Naive correction (y-axis) and similar statistics of xDF correction (x-axis) where, broadly speaking, large number of the connections with significant q-values no longer meet the 5% *α* level.

Since the changes in *Z*-scores due to xDF are spatially heterogeneous, both statistical and proportional thresholding methods are dramatically affected. In statistical thresholding (ST), after xDF correction, the FDR critical values (shown as dotted lines on Figure 6.A) are slightly increases from 2.064 to 2.14. With the use of xDF, 13.66% of the edges change from being marked FDR-significant to being non-significant; i.e. over 10% of the edges would be incorrectly selected with the Naive method. Similarly, proportional thresholding (PT) is also affected since the cost-efficient densities (shown as solid lines on Figure 6.A, see section S4.4 for more details on cost-efficient densities) are decreased from 35% to 27.5%. These changes in cost-efficient density result in 16.61% false positive edges meaning that they were found to be significant merely due to the autocorrelation effect.

The same analysis on another HCP subject finds very similar changes in functional connectivity (Figure S10) as in ST and PT, the critical values and cost-efficiencies were reduced by more than 50% and 26%, respectively, due to xDF correction. This results in 16% FP edges in ST and 19% FP edges in PT.

While Figure 6.A shows that there is a profound effect of xDF relative to no correction (Naive), it is of interest to see how xDF compares to an existing methods that does attempt to correct for autocorrelation. For this we compare xDF *Z*-scores to BH *Z*-scores (Figure 6.A); recall that BH correction does not account for cross-correlation Σ_*XY*_ and, due to confounding of cross-correlation and autocorrelation, can over-estimate *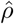* standard errors. When the *Z*-scores are low (corresponding to weak correlation) there is little difference between the approaches, while for stronger effects the difference between the two correction methods become clearer. For example, the *Z*-score for the edge between node 103 and node 104 (green dot in Figure 6.B; *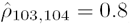*) with BH and xDF correction is 6.93 and 15.30, respectively; suggesting that the confounding in autocorrelation estimates (see Appendix D) has reduced the functional strength of this edge for more than 50%. Similarly, we also follow changes in rsFC of another HCP subject for nodes 23 and 88 (green dot in Figure S10.B; *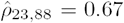*) where the confounding effect produces a similar effect; *Z*_BH_(23, 88) = 9.95, *Z*_xDF_(23, 88)=14.26.

Further, in Figure 6.D we show the impact of autocorrelation correction on mean value of rsFC *Z*-scores for edges included in proportional (left) and FDR-based statistical (right) thresholding. Reflecting the findings of the simulations, autocorrelation correction reduces *Z*-scores (suggesting Naive’s under estimation of *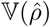*, but the BH method has appreciably smaller *Z*-scores (attributable to the statistical confounding problem discussed in Appendix D).

### 3.4 Effect of Autocorrelation Correction on Graph Theoretical Measures

Graph measures are notorious for their sensitivity to changes in functional connectivity (van den Heuvel et al., 2017). Using subjects from 100 HCP unrelated package, we show how accounting for autocorrelation can influence basic graph theoretical measures such as centrality and efficiency in weighted and binary networks. In weighted networks we use xDF-corrected standardised *Z*-scores as edge weights while in binary networks we set supra-threshold edges to one, zero otherwise.

The graph theoretical measures are discussed for both proportional (PT) and FDR-based statistical thresholding (ST) methods. In the former we use universal density (obtained from averaging cost-efficient densities across the three method under the investigation) to threshold rsFC across subjects while in latter we use hypothesis testing (*H*_0_ : *ρ*_*XY*_ = 0) for each pair of nodes. It is important to note that PT is a equi-density method while ST is equi-threshold method, meaning that the in proportionally thresholded matrices the number of edges is identical across all subjects and correction methods, while in statistically thresholded matrices the number of edges varies across subjects and correction method. For more information on each of the thresholding method, see Section S4.3 of the Supplementary Materials. We further repeat these analysis for the case of unthresholded rsFC, see Figure S11 for results of unthresholded functional connectomes.

Figure 7.A uses Bland-Altman plots to show the changes in graph measures of rsFC obtained using proportional thresholding. The left column of Figure 7.A shows the impact of xDF relative to no correction: Most dramatic is the overall reduction in weighted degree, with the degree hubs having the highest rate of losing weighted degrees; these nodes are parts of the Default Mode Network (DMN; i.e. DefaultABC) and Saliency Ventral Attention Network (SVAN; i.e. SalVenAtt). Similarly, the local efficiency of more than 98% of nodes were changed after xDF correction. Parts of DMN and SVAN in addition to parts of the Visual network are among the nodes which have been affected the most. In contrast, betweenness centrality has experienced modest changes of only *≈*5% in their values.

**Figure 7:**
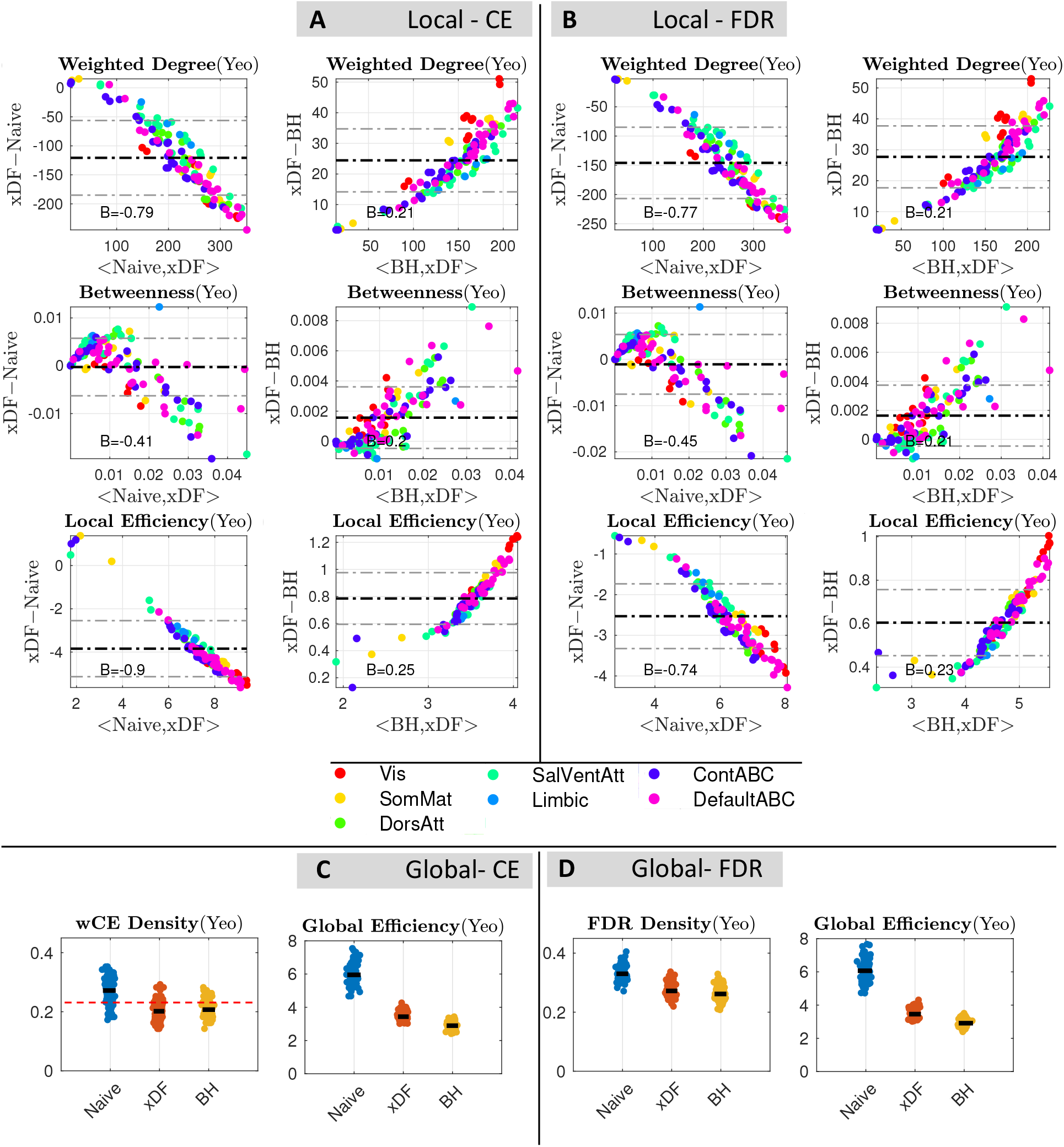
Overall changes in global and local graph theoretical measures with the 100 unrelated HCP package parcellated by Yeo atlas. **Panel A**, Bland Altman plots of xDF vs. Naive for weighted degree (top), betweenness (middle) and local efficiency (bottom) computed with a cost-efficient threshold. There is one point for each of 114 nodes, the particular measure averaged over subjects, and the nodes are colour coded according to their resting-state network assignment. **Panel B** Shows the same graph measures, but with statistical thresholding (corrected via FDR correction). **Panel C** shows the differences in weighted CE density (left) and Global efficiency (right), and **Panel D** illustrates the same results statistical FDR thresholding. There is a dramatic impact of correction method on all graph metrics considered. Similar results for Gordon (Figure S12), Power (Figure S13) and ICA200 (Figure S14) is available in Supplementary Materials.

The left column of Figure 7.B illustrates the changes in local graph measures of FDR-based statistical thresholded rsFC. Similar to ST results, the weighted degree of nodes suggests a significant reduction, with degree hubs (parts of DMN and SVAN) having the largest reduction. Local efficiency of almost every node (*≈* 99%) were affected, especially highly efficient nodes appear to be mostly influenced by xDF correction with parts of the DMN, SVAN and Visual network among them. Although the pattern of changes in betweenness centrality suggest almost no relation to the betweenness value of nodes before the correction, betweenness centrality of *≈* 22% of nodes are yet affected.

The right columns of Figures 7.A and 7.B reflect how the BH estimator is a more conservative correction (due to correlation-autocorrelation confounding effect; see Appendix D). Table 3 summaries changes due to the confounding effect; Visual network, DMN and SVAN are among the nodes which are most impacted.

**Table 3:** Percent of nodes that their weighted graph measures have significantly affected, for xDF vs. Naive and xDF vs. BH (see Figure 7). This quantifies the dramatic impact of correction method on each graph metric across all parcellation schemes and correction methods. For similar results for other parcellation schemes, see Table S2-S5.

| Comparison | Thresholding | Weighted Degree | Weighted Betweenness | Local Efficiency |
| --- | --- | --- | --- | --- |
| Naive > xDF | PT (CE) | 95.61% | 63.16% | 98.24% |
|  | ST (FDR) | 95.61% | 61.40% | 99.12% |
| BH > xDF | PT (CE) | 85.08% | 1% | 95.62% |
|  | ST (FDR) | 92.98% | 1.75% | 95.61% |

We also evaluate changes in the global measures. Figure 7.C shows the changes in network density (left) where the density of networks with Naively corrected network were significantly reduced after xDF and BH correction however there is a slight, yet significant, difference between density of xDF-corrected and BH-corrected networks. Similarly, the global efficiency of networks (Figure 7.C) are significantly reduced after accounting for autocorrelation. Figure 7.C also suggests that overestimation of variance due to correlation-autocorrelation confounding may yet reduce the local efficiency.

In the global measures a similar pattern of changes is also found for CE-based proportional thresholded networks, as the density (Figure 7.D) of xDF-corrected networks are significantly reduced. However, in spite of changes in weighted degree, the confounding effect may not affect the network densities. Finally, global efficiency of xDF-corrected networks suggests a significant reduction. Despite a small difference, the confounding effect has also reduced the global efficiency.

We repeat this analysis for the Gordon and the Power atlas. For the Power atlas, Figure S13 suggests that the autocorrelation leaves a very similar pattern of changes for both PT and ST. The highest changes take place in nodes with highest degree and efficiency measures; nodes comprising Visual, Fronto-parietal and Default Mode Network (DMN). For Gordon atlas (Figure S12), we found very similar results where the changes suggest that nodes from DMN, Fronto-parietal and Sensory-Motor (i.e. SMHand) networks experienced the highest changes in their weighted degree and local efficiency. Interestingly, similar to the Yeo atlas, Betweenness centrality has shown the highest resilience towards these changes. Finally, Figure S14 shows similar results for subjects parcellated using ICA200.

In Figure 8 we show the results of the comparison among Naive, xDF and BH repeated for *binary* graph measures derived from Yeo atlas. In absence of edge weights, any differences found are solely attributable to topological changes. The results suggest that the changes are still prominent between Naive and xDF for both proportionally and statistically thresholded rsFC maps, while there are no differences detected in graph metrics of binary networks corrected with xDF vs. BH. In Table 4 we summarise changes after correcting for autocorrelation with xDF and BH. These changes were tested across nodes between the two methods and corrected via FDR. For similar analysis with different parcellation schemes, see Figure S15 and S16.

**Table 4:** Percent of nodes that their binary graph measures have significantly affected, for xDF vs. Naive and xDF vs. BH (see Figure 8) in Yeo atlas. In xDF vs. Naive all graph measures were found to be significantly affected while in xDF vs. BH none of the measures suggest a change. For similar results for other parcellation schemes, see Table S6-S9

| Comparison | Thresholding | Binary Degree | Binary Betweenness | Local Efficiency |
| --- | --- | --- | --- | --- |
| Naive > xDF | PT (CE) | 95.61% | 63.16% | 98.24% |
|  | ST (FDR) | 95.61% | 48.24% | 28.07% |
| BH > xDF | PT (CE) | 85.087% | 0.877% | 95.61% |
|  | ST (FDR) | 0% | 0% | 0% |

**Figure 8:**
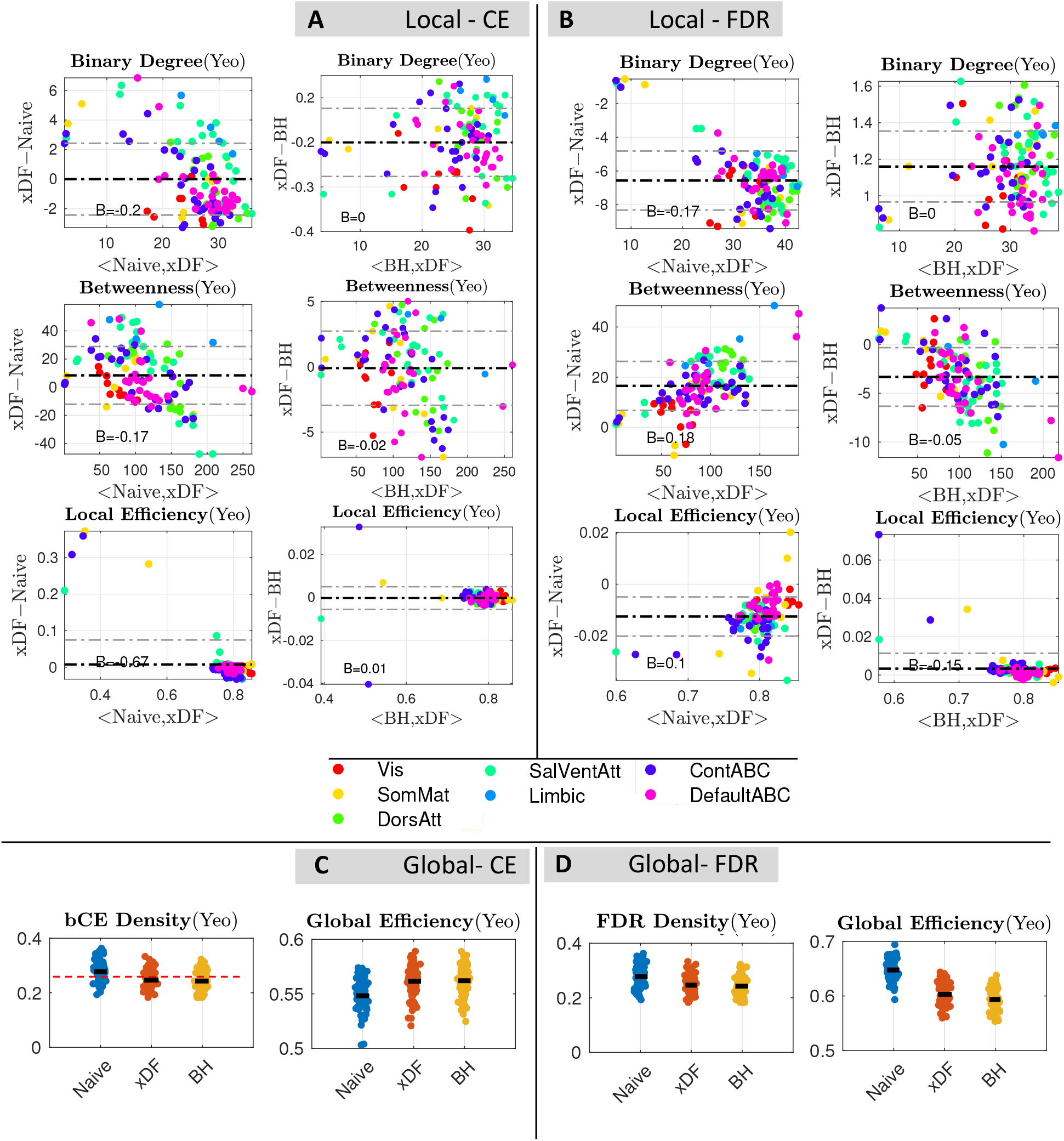
Overall changes in global and local graph theoretical measures with the 100 unrelated HCP package parcellated by Yeo atlas. **Panel A**, Bland Altman plots of xDF vs. Naive for binary degree (top), betweenness (middle) and local efficiency (bottom) computed with a cost-efficient threshold. There is one point for each of 114 nodes, the particular measure averaged over subjects, and the nodes are colour coded according to their resting-state network assignment. **Panel B** Shows the same graph measures, but with statistical thresholding (corrected via FDR correction). **Panel C** shows the differences in weighted CE density (left) and Global efficiency (right), and **Panel D** illustrates the same results statistical FDR thresholding. There is a dramatic impact of correction method on all graph metrics considered. Similar results for Gordon (Figure S15), Power (Figure S16) and ICA200 (Figure S17) is available in Supplementary Materials.

## 4. Discussion

We have developed an improved estimator of the variance of the Pearson’s correlation coefficient, xDF, that accounts for the impact of autocorrelation in each variable pair as well as the instantaneous and lagged cross-correlation. On the basis of extensive simulations under the null setting (*ρ* = 0) using simulated data and real data with inter-subject scrambling, the xDF, BH and Q47 methods have good control of false positives, with xDF showing only slight FPR inflation on real null data (5.7%) and, on simulated data, only inflated with strong autocorrelation for short time series. Naive (no correction) has severe inflation of FPR as do other methods based on simplistic AR(1) autocorrelation models (G-Q47, AR1MCPS) or common ACF for each pair of variables have poor FPR control. Simulations with realistic autocorrelation and non-null cross-correlation find that Naive severely under-estimates variance while BH and Q47 over-estimates variance, likely due to a confounding of auto-and cross-correlation in those corrections; xDF, in contrast, has negligible bias for long time series and for short time series has low bias for all but the strongest forms for autocorrelation.

On real data (non-null) rsFC we replicate the simulation findings, with Naive *Z*-scores dramatically inflated relative to xDF, BH and Q47 *Z*-scores smaller in magnitude. The differences between the methods, however, are node specific, reflecting how xDF adjusts for autocorrelation in each node pair. We recommend that all rsFC analyses that are based on *Z*-scores, whether thresholded arbitrarily or or, say, use a mixture modelling approach (Bielczyk et al., 2018), use the xDF correction to obtain the most accurate inferences possible.

We show that graph analysis measures are dramatically impacted by use of xDF, relative to either Naive or BH corrections. Broadly speaking, accounting for autocorrelation results in lower *Z*-scores and lower rsFC densities. These heterogeneous changes alter the topological features of the functional connectome, however the changes are not similar across resting-state networks; in the HCP data, we find nodal strengths and local efficiencies in parts of the subcortical regions experience lower changes compared to nodes from the frontoparietal and default mode networks, which are among the highly affected areas. The pattern of changes suggest that the nodal degree and efficiency hubs are among the most affected. In contrast, results for betweenness centrality suggest no systematic pattern with relatively lower changes.

We provide a comprehensive review of the literature on autocorrelation corrections for the variance of the sample correlation, usually cast as estimation of the effective degrees of freedom. In the neuroimaging community this is sometimes referenced as “Bartlett Correction Factor (BCF)”, though it has been only informally defined and used as a global correction over subjects and ROIs (Hale et al., 2016). We emphasize the importance of truncation or tapering of ACF’s and computing a correction for each node pair.

We note the strong influence of ROI size on the strength of autocorrelation, with, at one extreme, voxel-level data having the weakest correlation, increasing in strength as the size of the ROI increases; an effect that is often ignored in rsFC studies (Lee and Xue, 2017) or indirectly by regressing out the volume of the ROIs (Sethi et al., 2017). We also showed that, even for an ROI atlas with identically sized regions (i.e. Power atlas), autocorrelation can vary substantially over the brain depending on their location. These factors, along with inter-subject heterogeneity of the autocorrelation effect (Figure 2.E), could become a significant source of bias in any rsFC analysis using *Z*-scores if not otherwise corrected. And we note that our findings hold for both volumetric and surface-based analysis (Figure 2.C).

We stress that our work does not invalidate use of correlation entirely, as our derivation shows that Pearson’s correlation is approximately unbiased for the correlation *ρ* in the data (Appendix B). For between-subject analyses, the varying intra-subject standard deviation of Pearson’s correlations is analogous to fMRI, where some methods ignore first-level standard errors, which has been shown to be valid for simple models (Mumford and Nichols, 2009) (e.g. in SPM and AFNI’s 3dDeconvolve), while other methods account for these standard errors (FSL’s FLAME and AFNI’s 3dMEMA). For group level inference on correlations, we are only aware of the work of Fiecas et al. (2017) that provides an analogous 2-level model that accounts for intra-subject standard errors. One way forward is a hybrid approach, where any thresholding is done on xDF *Z*-scores, and then subsequent analyses are done on surviving *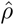* values (see, e.g. Bassett et al. (2011), that uses such an approach but with Naive Z).

In short, any rsFC analysis based on *Z*-scores must ensure that the calculation of those *Z*-scores account for the impact of temporal autocorrelation in a subject-and edge-specific manner, as with our xDF method.

As an aside, we note that our statement on the unbiasedness of correlation is at odds with other recent work (Arbabshirani et al., 2014; Davey et al., 2013). This is not inconsistent: Both of these works start their analysis with a pair of white noise variables with instantaneous correlation *ρ* and then assess the impact of inducing autocorrelation on those variables. In particular, they note that if different autocorrelation structures are induced then a bias in estimate of *ρ* can arise. Instead, we study the auto-and cross-correlation of the presented data (*X, Y*), without reference to an (unobserved) autocorrelation-free signal; in this setting Pearson’s correlation is (approximately) unbiased regardless of differential autocorrelation. We believe our empirical approach is more appropriate, as the BOLD signal is not white and hence inference on the correlation of presented (and not latent white) signals is of primary interest.

We note that other authors have proposed pre-whitening as a solution to improve inference on correlation (Bright et al., 2017), and that pre-whitening is recommended when conducting system identification of the cross-correlation function *ρ*_*XY,k*_ (Priestley, *1983). However, we still see the value of the no-whitening plus xDF correction approach. First, pre-whitening requires accurate estimation of Σ*_*X*_ and Σ_*Y*_, and careful evaluation is required to see if the FPR is controlled and the standard error unbiased over a range of settings. Second, pre-whitening changes the definition of *ρ*, from concerning the instantaneous correlation of the observed time series to that of the (unobserved, latent) white time series. And, perhaps most important for sliding window time-varying rsFC, pre-whitening mixes data from distant time points with neighbouring ones, challenging the interpretation of the individual time points as pertaining to a precise moment in time. Pre-whitening is also used with voxelwise linear modeling using a seed region predictor. This is again different, in that the same whitening – based on voxelwise residuals – is applied to both response and predictor, perhaps improving the interpretability of this approach over the case of separate whitening for each *X* and *Y*. It is difficult to predict how this approach will differ from correlation inference with xDF, as xDF considers autocorrelation of both time series as well as cross-correlation, while this approach only considers voxelwise residual autocorrelation.

One immediate extension to the current work is to adapt the xDF estimator to partial correlations. Partial correlations have recently drawn substantial attention after they were shown to be effective in resting-state analysis (Marrelec et al., 2006; Smith et al., 2011). Further, recent studies have shown that accounting for autocorrelation is remarkably sensitive to sampling rates of the fMRI BOLD time series (Bollmann et al., 2018), therefore evaluating the proposed methods on different sampling rates would be useful. We have not attempted to investigate how the changes in *Z*-scores we describe would affect the inter-group changes, but this would be a useful extension, as Váša et al. (2018) did for Schizophrenia in context of wavelet EDF. Finally, it is important to note that application of the xDF is not confined to rsFC of fMRI time series as it can be used in other modalities such as EEG and MEG as both modalities were shown to suffer from dependencies amongst their data-points.

## 5. Software Availability and Reproducibility

Analysis presented in this paper have been done in MATLAB 2015b and R 3.1.0. Graph theoretical analysis were done using Brain Connectivity Toolbox (accessed: 15/1/2017) (Rubinov and Sporns, 2010).

Variance of Pearson’s correlation, *Z*-scores and p-values of such correlation matrices can be estimated via `xDF.m` available in https://github.com/asoroosh/xDF. The script is an standalone function and is executable using Statistics and Machine Learning Toolbox in MATLAB 2016 or later. The repository also contains six other variance estimators discussed in this work.

The autocorrelation (`AC fft.m`) and cross-correlation (`xC fft.m`) functions are estimated using Wiener–Khinchin theorem which involves discrete Fourier transformation of time series. We also used an algorithm proposed by Higham (1988) to find the nearest positive semi-definite covariance matrices for simulations described in Section S3.

Scripts and instructions to reproduce all the figures and results, are also available via http://www.github.com/asoroosh/xDF_Paper18. For details regarding the reproduciblity of the figures see section S6 of the supplementary materials.

## 6. Acknowledgements

We thank Andrew Zalesky at University of Melbourne, Mark Fiecas at University of Minnesota, Simon Schwab and Samuel Davenport at University of Oxford and Jeanette Mumford at University of Wisconsin-Madison for their useful input.

T.E.N. is supported by the Wellcome Trust (100309/Z/12/Z). S.M.S. is supported by the Wellcome Trust Centre for Integrative Neuroimaging (203139/Z/16/Z) and the Wellcome Trust Strategic Award “Integrated Multimodal Brain Imaging for Neuroscience Research and Clinical Practice” (098369/Z/12/Z).

Data were provided by the Human Connectome Project, WU-Minn Consortium (Principal Investigators: David Van Essen and Kamil Ugurbil; 1U54MH091657) funded by the 16 NIH Institutes and Centers that support the NIH Blueprint for Neuroscience Research; and by the McDonnell Center for Systems Neuroscience at Washington University.

Computation used the BMRC facility, a joint development between the Wellcome Centre for Human Genetics and the Big Data Institute supported by the NIHR Oxford BRC. The views expressed are those of the authors and not necessarily those of the NHS, the NIHR or the Department of Health.

## A. Results for Joint Distribution of Time Series *X* and *Y*

Here we provide basic results required for the next appendix, for moments of inner and cross produces of *X* and *Y*.

### Theorem 1

(Covariance of Quadratic Form of Bivariate Gaussian Distribution). *For fixed matrices* **A** *and* **B**, *if G is a random vector such that G ∼ N* (0, **Φ**) *then* ℂ(*G*^┬^**A***G, G*^┬^**B***G*) = 2 tr(**ΦAΦB**)

*Proof*. The result follows from application of the definition of covariance,

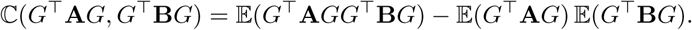

and expectation of a quadratic form for Gaussian variates (Petersen et al., 2008),

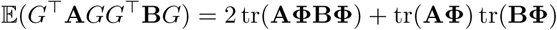

and expectation of quadratic forms, 𝔼(*G*^┬^**A***G*,) = tr(**A Φ)** and 𝔼(*G*^┬^**B***G*,) = tr(**B Φ)**. □

The Gaussian assumption can be relaxed, but then an additional term arises to account for departures from Gaussian kurtosis.

The inner products of *X* and *Y* can now be represented in terms of a quadratic form of

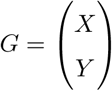

for following maticies:

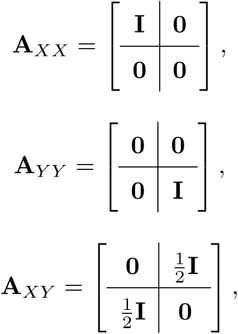

such that *X*^┬^*X* = *G*^┬^**A**_*XX*_*G, Y* ^┬^*Y* = *G*^┬^**A**_*Y Y*_ *G*, and *X*^┬^*Y* = *G*^┬^**A**_*XY*_ *G*.

## B. xDF: Variance of Sample Correlation Coefficient for Arbitrary Dependence

For mean zero length-*N* random vectors *X* and *X* with (*N × N*) variance matrices Σ_*X*_ and Σ_*Y*_ and cross-covariance Σ_*XY*_, we develop the variance of the sample correlation. Following Lehmann (1999a) and Hunter (2014), we derive an approximation for the sampling variance of Pearson’s correlation. Starting with the 3-dimensional sufficient statistic

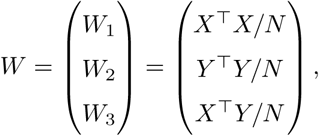

note that the function 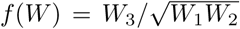 generates the correlation coefficient *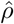*. Then the first order Taylor’s series of *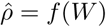* about 𝔼(*W*) is

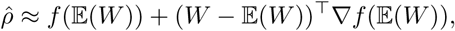

so that 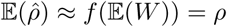 and

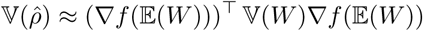

where the gradient of *f* (·) is

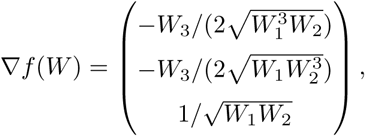

evaluated at *W* = E(*W*),

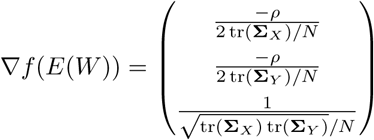

and, by Theorem 1 in Appendix A,

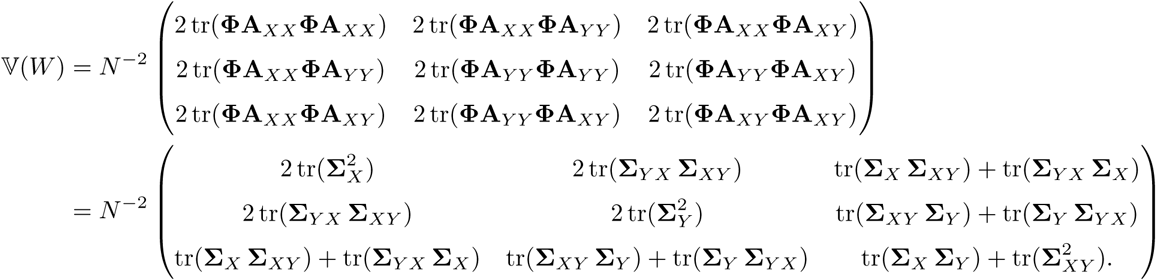

Based on these expressions, evaluating the matrix product (*∇f* (𝔼(*W*)))^┬^ *𝕍*(*W*)*∇f* (𝔼(*W*)) gives the result in Eq. 1. While very similar, this derivation is not a standard delta method result as we do not have independent observations.

## C. Trace of product of two Toeplitz Matrices

If *ρ*_*XX,k*_ and *ρ*_*Y Y,k*_ are autocorrelation coefficients of time series X and Y on lag *k*, the diag(**Σ**_*X*_ **Σ**_*Y*_) can be re-written as

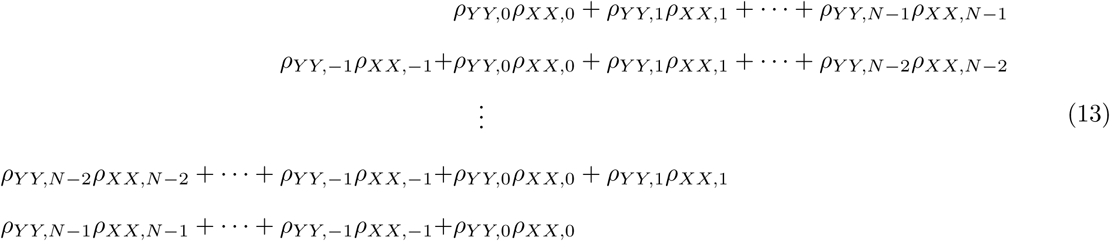

Considering that the autocorrelation function of time series is symmetric (i.e. the negative and the positive lags are identical), for X and Y we can simply obtain the trace of the **Σ**_*X*_ **Σ**_*Y*_ as,

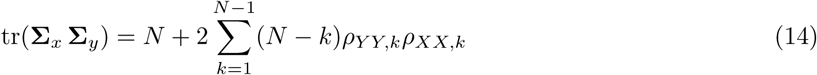

Similarly, the covariance matrix, **Σ**_*XY*_, can be written as a Toeplitz matrix of form

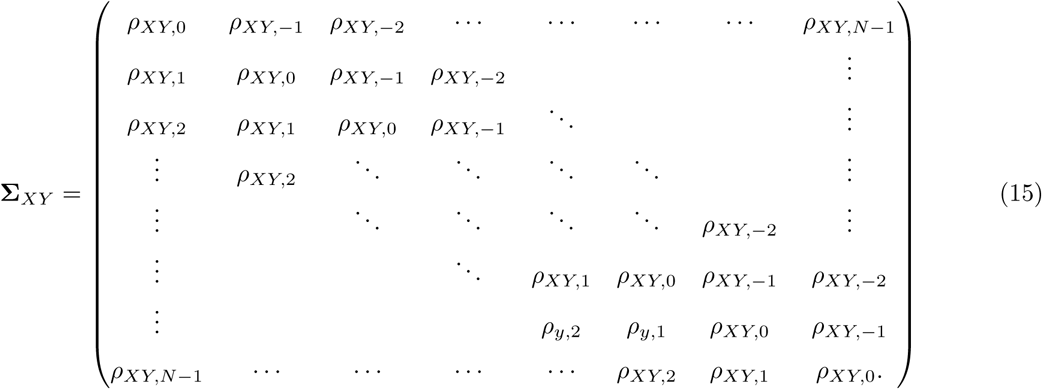

Knowing that the cross-covariance matrices are asymmetric but 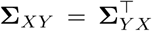, the trace of tr 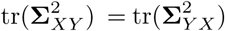 and can be re-written as

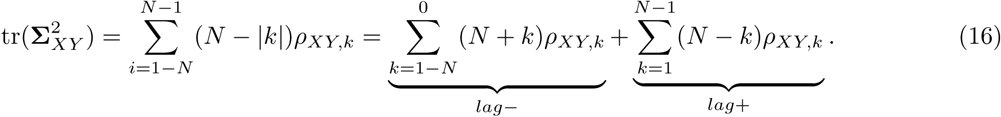

Similarly, other terms of the Eq. 1 can be re-written as vector operations (see Eq. 2).

## D. Confounding of Autocorrelation and Cross-correlation Estimates

A majority of the EDF estimators discussed have the term tr(**Σ**_*X*_ **Σ**_*Y*_) which depends on the product of autocorrelation functions, *ρ*_*XX,k*_*ρ*_*Y Y,k*_ (see Appendix C). Some unexpected results such as poor variance estimation for the seemingly easy case of time series with no autocorrelation (see Section 3.2) were found to be caused by confounding between estimates of autocorrelation and cross-correlation. Observe that for two white but dependent time series (*ρ*_*XX,k*_ = *ρ*_*Y Y,k*_ = 0 for *k >* 0, but *ρ*_*XY*,0_ ≠0), the product of the sample autocorrelation functions are

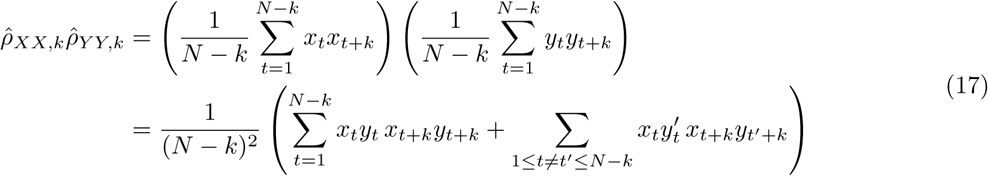

In this last expression note that each term in the first sum has an expected value of *ρ*^2^, while the second sum has an expected value of zero. As a result, even when there is no (or very light) autocorrelation, methods dependent on tr(**Σ**_*X*_ **Σ**_*Y*_) or *ρ*_*XX,k*_*ρ*_*Y Y,k*_ alone can be substantially biased by non-zero cross-correlation. This includes BH and all its variations listed in Pyper and Peterman (1998). Our xDF, on the other hand, is immune of the effect thanks to the cancelling cross-covariance terms in Eq. 2.

## E. Autocorrelation in Regions of Interests

We observed substantial differences in severity of autocorrelation in voxel-wise data as opposed to data derived from regions of interest (ROI); see Fig. 2. This section proposes a model that explains the effect.

Suppose that a ROI contains *R* voxels, and the data for time *t* and voxel *i* = 1, …, *R* can be be modelled as,

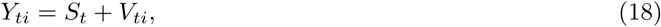

where the *S*_*t*_ a common “signal” shared across all the voxels within the ROI (i.e. a ROI-specific global signal) and *V*_*it*_ are the voxel-specific component of data. Assuming that the two components are uncorrelated, the autocorrelation for the ROI is defined as,

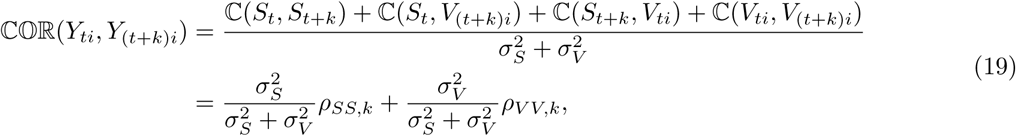

where 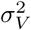 and 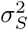 are the variances of *V* and *S*, respectively, and *ρ*_*V V,k*_ and *ρ*_*SS,k*_ are the autocorrelation coefficient of *V* and *S* at lag *k*, specifically. This shows that the voxel-level correlation is a convex combination of the two ACFs, balanced according to the variances 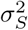 and 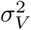.

Now, if we assume for this illustration that voxels are independent, then for the ROI-averaged time series, 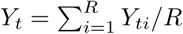, the autocorrelation is:

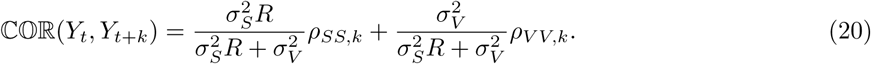

We again have a convex combination, but now balanced according to 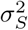 *R* and 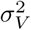.

If we make some reasonable assumptions we can make predictions of this correlation as *R* grows. Let us assume that *ρ*_*SS*_ reflects a stronger autocorrelation of a (BOLD-related) common signal, and *ρ*_*V V*_ reflects less autocorrelation and more thermal noise contributions at individual voxels. Then as *R* grows the ACF of the ROI average converges to the ACF of the common signal, and this would explain the increased strength of autocorrelation with larger ROIs.

## Footnotes

1 While not the topic of this work, Arbabshirani et al. (2014) also discuss bias in sample correlation coefficients 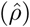 due to autocorrelation. In contrast, our derivation (Appendix B) finds no such bias; see Section 4.

2 http://fsl.fmrib.ox.ac.uk/fsl/fslwiki/FSLNets; visited on 18 September 2015

