## Supplementary Materials for "Effective Degrees of Freedom of the Pearson’s Correlation Coefficient under Autocorrelation"

#### S1 Introduction

In first part of this document we start with describing the background to correlation coefficients and correlation matrices as measure of functional connectivity, we later explain the validations methods we used to compare the methods explained in Section 3.2. We then present a detailed description of the resting-state data and parcellation schemes we used in this study. Further, we discuss conventional thresholding techniques (i.e. statistical, proportional) and graph theoretical measures we employ to demonstrate the impact of xDF correction.

In the second part, we discuss the results of Oracle simulation aiming to verify the xDF. We then use Monte Carlo simulations to evaluate the xDF performance across different range of tapering methods. Finally, we discuss the impact of xDF correction on functional connectivity as well as graph theoretical measures for two more parcellation atlases, Power and Gordon, aiming to ensure the robustness of the results presented in this work.

#### S2 Background

To form a functional connectivity of each subject, we use Pearson's correlation to measure the temporal similarity between each pair of the time series, $x$ and $y$, representing a brain region. Therefore, each ROI is considered as a node and the correlation coefficient between $X$ and $Y$ is considered as the weight to the edge formed between the two nodes. The population ('true') correlation coefficient, $\rho_{XY}$, between two random variables $X$ and $Y$ is

$$\rho_{XY} = \frac{\mathbb{C}(X,Y)}{\sqrt{\mathbb{V}(X)\,\mathbb{V}(Y)}}. \tag{S1}$$

For samples of $X$, $x_1, x_2, ..., x_N$, and of $Y$, $y_1, y_2, ..., y_N$, the Pearson's correlation coefficient is an estimate of $\rho_{XY}$,

$$\hat{\rho}_{XY} = \frac{\sum_{i=1}^{N}(x_i - \bar{x})(y_i - \bar{y})}{\sqrt{\sum_{i=1}^{N}(x_i - \bar{x})^2}\sqrt{\sum_{i=1}^{N}(y_i - \bar{y})^2}}, \tag{S2}$$

where $N$, is the number of observations and $\bar{x}$ and $\bar{y}$ are the sample means of $X$ and $Y$, respectively. This estimate is only approximately unbiased (Kendall et al., 1994), and for the case of N independent pairs of samples

$$\mathbb{E}(\hat{\rho}_{XY}) \approx \rho_{XY} - \frac{\rho_{XY}(1-\rho_{XY}^2)}{2N}, \tag{S3}$$

and has variance that varies with the true correlation (Kendall et al., 1994)

$$\mathbb{V}(\hat{\rho}_{XY}) \approx \frac{(1-\rho_{XY}^2)^2}{N}\left(1 + \frac{11\rho_{XY}^2}{2N}\right). \tag{S4}$$

In addition to the variance being dependent on the true $\rho_{XY}$, the sampling distribution can be highly non-normal. To address this, Fisher's transformation is typically applied as following

$$\hat{z}_{XY} = \text{arctanh}(\hat{\rho}_{XY,0}) = \frac{1}{2}\ln\left[\frac{1+\hat{\rho}_{XY,0}}{1-\hat{\rho}_{XY,0}}\right]. \tag{S5}$$

When the original data comprise independent, identical draws from a bivariate normal distribution, Fisher's $z_{XY}$ has an approximate normal distribution with mean acrtanh($\rho_{XY}$) and variance $1/(N-3)$. Thus a standardised statistic, a Z-score, can be computed as $Z = \hat{F}_{XY}\sqrt{N-3}$.

### S3 Validation Methods

#### S3.1 Simulation Settings

Without loss of generality, we seek to simulate mean zero and unit variance time series $X$ and $Y$, each of length $N$, such that $\mathbb{V}(X) = \Sigma_X$, $\mathbb{V}(Y) = \Sigma_Y$ and lag-0 correlation $\mathbb{COR}(X_i, Y_i) = \rho_{XY}$. The stacked time series of $(2N)$-vector, $\mathbf{G} = \begin{pmatrix} X \\ Y \end{pmatrix}$, can be simulated from a (2N)-vector $\mathbf{W}$, with $\mathbb{E}(W) = 0$ and $\mathbb{V}(\mathbf{W}) = \mathbf{I}$, as

$$\mathbf{G} = \mathbf{B}\mathbf{W} \tag{S6}$$

with the $2N \times 2N$ matrix $\mathbf{B}$ is defined as

$$\mathbf{B} = \begin{pmatrix} a_{XX}\mathbf{K}_X & a_{XY}\mathbf{K}_X \\ a_{YX}\mathbf{K}_Y & a_{YY}\mathbf{K}_Y \end{pmatrix}, \tag{S7}$$

where $\mathbf{K}_X$ is chosen such that $\mathbf{K}_X\mathbf{K}_X^\top = \mathbf{\Sigma}_X$, $\mathbf{K}_Y$ such that $\mathbf{K}_Y\mathbf{K}_Y^\top = \mathbf{\Sigma}_Y$, and $\mathbf{A} = \begin{pmatrix} a_{XX} & a_{XY} \\ a_{YX} & a_{YY} \end{pmatrix}$ is chosen such that $\mathbf{A}\mathbf{A}^\top = \begin{pmatrix} 1 & \rho^*_{XY} \\ \rho^*_{XY} & 1 \end{pmatrix}$. The simulated data $\mathbf{G}$ will then have

$$\mathbb{V}(G) = \begin{pmatrix} \mathbf{\Sigma}_X \, \mathbf{\Sigma}_{XY} \\ \mathbf{\Sigma}_{XY}^\top \, \mathbf{\Sigma}_Y \end{pmatrix}, \tag{S8}$$

where $\mathbf{\Sigma}_{XY} = \rho^*_{XY}\mathbf{K}_X\mathbf{K}_Y^\top$ is the induced cross correlation. While edge effects will mean $\mathbf{\Sigma}_{XY}$ doesn't have a constant diagonal, we can obtain the desired instantaneous cross-correlation $\rho_{XY}$ by setting

$$\rho^*_{XY} = \frac{\rho_{XY}}{\mathrm{tr}(\mathbf{K}_X\mathbf{K}_Y^\top)/N}. \tag{S9}$$

We simulate time series pair of length $N = \{100, 200, 600, 1200\}$ with correlation coefficients $\rho = \{0, 0.2, 0.5, 0.7, 0.9\}$ between the each time series. Six different autocorrelation structure were used, listed in Table S1. Although studies are commonly use synthetic autocorrelation structure, in this work we use autocorrelation of real world data-sets.

Table S1: Autocorrelation structures used in Monte Carlo Simulations.

| Labels | AR lags | Source | Adjustments |
|---|---|---|---|
| W | 0 | N/A | N/A |
| AR1 | 1 | N/A | N/A |
| AR4 | 4 | Germany Population Growth Rate | N/A |
| AR14 | 14 | ROI 35 of HCP Subject 135923 | FPP |
| AR20 | 20 | ROI 35 of HCP Subject 135923 | FPP |

This result in 300 unique combinations of autocorrelations, correlation coefficients and sample size. The pair time series of each combination is generated using the model explained in section S3.1 for 5K realisations. The bias of each method is calculated via Eq. S10,

$$\%Bias = \frac{Statistics - GroundTruth}{GroundTruth} \times 100 \tag{S10}$$

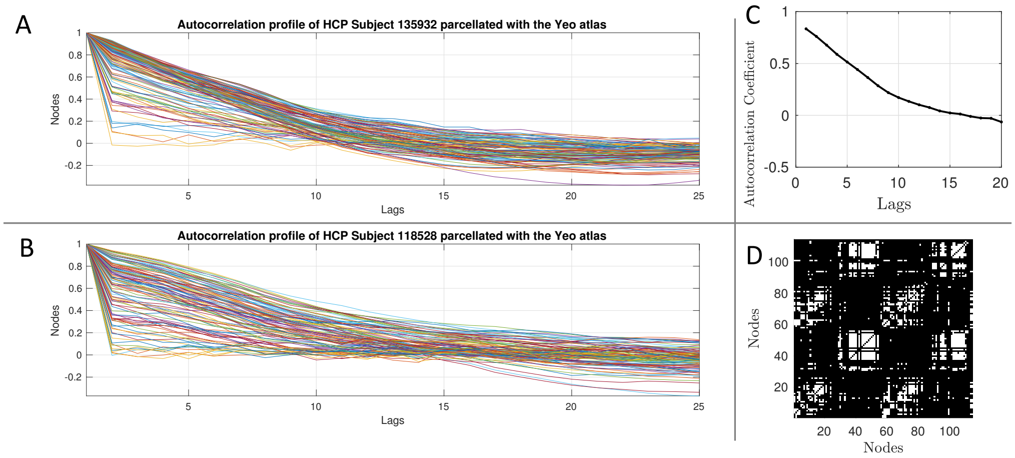


Figure S1: Autocorrelation structures used in Monte Carlo and Oracle simulations. **Panel A** illustrates the autocorrelation coefficients for the first 200 lags in Subject 135932 **Panel B** shows autocorrelation coefficients for Subject 118528 **Panel C** shows the first 20 lags of Node 35 in Subject 135932 **Panel D** is an example adjacency matrix of Subject 118528 on 15% proportional density.

#### S3.2 Oracle Simulations

Estimation of autocorrelation coefficients are notoriously noisy. Therefore, in addition to Monte Carlo simulation (discussed later in Section S3.3), we use oracle simulations to evaluate performance of each method under noise-free conditions where the autocorrelation are rather *calculated* than estimate. Algorithm 1 demonstrates an example for two time series $X$ and $Y$ of length $N = 600$ with AR4 and AR14 autocorrelation structure, respectively and correlation coefficient of 0.6. In terms of bias calculation, Eq. S10, the ground truth is the variance of correlation coefficients generated via MC simulations, $\sigma^2_{MC}$, and the Statistics is the calculated theoretical variance, $\sigma_O^2$.

**Algorithm S1:** Oracle Simulation Example

initialization;

$I \leftarrow 5k, \quad T \leftarrow 600,$ ;

$\rho_{XY,0} \leftarrow 0.6, \quad \rho_{XX} \leftarrow$ AR4, $\quad \rho_{YY} \leftarrow$ AR14

**while** $i < I, \quad i++$ , **do**

  Generate $X$ and $Y$ using Eq S6 ;

  $\hat{\rho}_{XY}(i) \leftarrow \mathbb{COR}(X, Y)$;

**end**

Estimate $\mathbb{V}(\rho_{XY})$ using $\rho_{XY,0}$, $\rho_{XY}$, $\rho_{YY}$, $\rho_{XX}$ via Eq. 1 of main text ;

$\sigma_{MC}^2 \leftarrow \mathbb{V}(\hat{\rho}_{XY})$;

$\sigma_O^2 \leftarrow \mathbb{V}(\rho_{XY})$;

%Bias $\leftarrow (\sigma_O^2 - \sigma_{MC}^2)/\sigma_{MC}^2 \times 100$  (See Eq. S10)

#### S3.3 Monte Carlo Simulation

We further, use Monte Carlo simulations to compare performance of xDF against the methods discussed in Section 2.2.1 and 2.2.2. We also use the same simulations to evaluate the tapering methods. Algorithm 2 demonstrate the procedure for an example where the sample size is 600, the correlation coefficients between two time series is 0.6 and the autocorrelation structures of two time series are AR4 and AR20-Yeo35, respectively. We finally evaluate each regularisation methods explained in Section 2.3 by repeating the simulations for each technique. Similar to Oracle simulations, the bias is calculated via Eq. S10 where the ground truth is the variance of correlation coefficients generated via MC simulations, $\sigma_{MC}^2$, but the Statistics is the mean of estimated variance via xDF, $\sigma_T^2$.

**Algorithm S2:** Monte Carlo Simulation Example

initialization;

$I \leftarrow 5k, \quad T \leftarrow 600,$ ;

$\rho_{XY,0} \leftarrow 0.6, \quad \rho_{XX} \leftarrow$ AR4, $\quad \rho_{YY} \leftarrow$ AR14

**while** $i < I, \quad i++$ , **do**

  Generate $X$ and $Y$ using Eq S6 ;

  $\hat{\rho}_{XX} \leftarrow$ autocorrelation $(X)$;

  update $\hat{\rho}_{XX}$ with a regularised version (see section 2.3 of main text);

  $\hat{\rho}_{YY} \leftarrow$ autocorrelation $(Y)$;

  update $\hat{\rho}_{YY}$ with a regularised version;

  $\hat{\rho}_{XY} \leftarrow$ cross-correlation $(X,Y)$;

  update $\hat{\rho}_{XY}$ with a regularised version;

  $\hat{\rho}_{XY,0}(i) \leftarrow \hat{\rho}_{XY}(0)$ ;

  $\mathbf{V}(i) \leftarrow$ Estimate $\mathbb{V}(\hat{\rho}_{XY,0})$ using $\hat{\rho}_{XY0}$, $\hat{\rho}_{XY}$, $\hat{\rho}_{YY}$, $\hat{\rho}_{XX}$ via Eq. 1 of main text ;

**end**

$\sigma^2_{MC} \leftarrow \mathbb{V}(\hat{\rho}_{XY,0})$;

$\sigma^2_T \leftarrow \frac{1}{I}\sum_i^I \mathbf{V}(i)$ ;

%Bias $\leftarrow (\sigma^2_T - \sigma^2_{MC})/\sigma^2_{MC} \times 100$ (See Eq. S10)

#### S3.4 False Positive Rates

In this section we compare the False Positive Rates (FPR) of correction methods, on 5,000 pairs of uncorrelated time series for each of the combinations explained in Section S3.1. For a desired $\alpha-$level, the FPR is calculated as ratio of significant Z-scores to number of realisations. Results of such analysis is presented in Figure 3.C of the main text.

#### S3.5 Correlation Matrices

Besides pair-wise Oracle and Monte Carlo simulations, we also use simulated correlation matrices to investigate the effect of global correction methods (i.e G-46 and AR1MCPS). The autocorrelation structure for each node in the correlation matrices is inherited from full-lagged autocorrelations of the time series in HCP subject 118528, parcellated with Yeo atlas (i.e. 114 nodes). Therefore, contrary to Monte Carlo simulations, sample sizes are limited to $T = 1200$. We conduct two independent analysis using such correlation matrices;

- Similar to pair-wise FPR analysis discussed in Section S3.4, we generate 2,000 simulated correlation matrix of *uncorrelated* time series. The FPR for three $\alpha-$levels=$\{1\%, 5\%, 10\%\}$ is then calculated as ratio of significant edges to number of total edges for a desired $\alpha$-level. We also evaluate how each correction method has affected the Z-scores using QQ-plots and Kolmogrove Smirnov (KS) statistics. Figure 3.B illustrates results of such analysis.

- Although uncorrelated time series, discussed in the previous item, are commonly used to evaluate the DF correction methods, they often ignore the extent of the aliasing effect. For this reason, we generate 2,000 correlation matrices, which 15% of edges within each matrix are correlated; correlation coefficients range from 0.2 and 0.9. We then quantify performance of each method by calculating the sensitivity, specificity and accuracy of each method.

##### S3.6 Inter-Subject Scrambling (ISS)

Due to non-reproducible nature of BOLD signals, the correlation coefficient between two randomly picked time series from subject $u$ and $q$, $\hat{\rho}_{uq}$ are expected to follow mean-zero unit variance Normal distribution, $\mathcal{N}(0,1)$. Therefore, after many realisations, any systematic confounds on subjects, such as autocorrelation, should inflates the variance of resultant correlation coefficients. A similar method has been used to find subject-level thresholding of functional connectivity (Smith et al., 2011; Bielczyk et al., 2018).

For each estimator, we repeat the inter-subject scrambling for 82k randomly picked nodes across 100 HCP subjects. We then use two-sided unequal Kolmogorov-Smirnov statistics, $L$, which is defined as the maximum of absolute difference between Cumulative Distribution Function (CDF) of the surrogate data, $\Psi$, and a mean-zeroed univariate normal distribution, $\mathcal{N}$, and can be formulated as below:

$$L = \max_{x}(|\Psi(x) - \mathcal{N}(x)|) \tag{S11}$$

Hence, value of $L$ closer to zero indicate the shorter the distance between the empirical and normal distributions. We also use QQ-plots and histograms to visually inspect the deviation of each estimator from the normal distribution.

#### S4 Real Data Analysis

We use 100 resting-state scans of standard Unrelated package in the HCP cohort where participants were asked to remain awake in a MRI machine with their eyes open. Each subject was asked to fixate on a projected bright cross-hair on a dark background. Gradient-echo EPI (TE=33.1ms, flip angle=52, FOV=208x180mm) was used to acquire a 15min scan of each subject with TR=720ms, which result in timeseries of 1200 datapoints for each subjects. Each brain volume were parcellated isomorphic voxels of $2mm^3$.

We used an extended pre-processed version of the data available via HCP repository. Each scan has undergone a thorough pre-processing procedure as explained in Glasser et al. (2013). Further to the traditional pre-processings steps, high-pass filtered data were denoised by the non-aggressive FIX algorithm (Salimi-Khorshidi et al., 2014) to ensure any 'bad' components, such as movement, physiological noises of heart beat and respiration, have been regressed out of each individual scan. As of the recent effort to show the spurious effect of the global mean signal (Ciric et al., 2017; Burgess et al., 2016; Power et al., 2017) we regress out the global mean in all data sets. The HCP Subject 101107 was removed prior to analysis due to significant head motions which results in missing data on orbitofrontal cortex. Quality of resting-state fMRI scans of the HCP 100 unrelated package has been discussed in details in Afyouni and Nichols (2018).

##### S4.1 Parcellation Schemes

Here we briefly discuss the parcellation schemes used in this work;

- **Yeo** is a data-driven atlas which was formed out of similar pattern of functional connectivity of voxels across two resting-state scans of 500-subjects cohorts of healthy brains. Networks of functionally coupled regions were identified by employing a machine-learning clustering method (Yeo et al., 2011). Resultant ROIs are dropped into 7 Resting-state networks (RSN): Default Mode Network, Dorsal Attention Network, Saliency Ventral Attention Network, Control Network, Limbic regions, Visual and Somatomotor Networks. However, these 17 RSNs, are also subdivided into 17 networks. These 17 networks were found across a surface-based realignment of all subjects which merely allows investigating the cortical regions of the human brain. This parcellation scheme delivers a clear evidences of segregation/integration nature of the human brain function as well as hierarchical structures of the connectivity profiles.

- **ICA200** is data-driven atlas which was formed by applying Independent Component Analysis (ICA) on resting-state functional scans of 1200 human brain. ICA is a mathematical method of distinguishing between the latent number of additive components of a multivariate signal (Beckmann and Smith, 2004). Applying ICA on the brain fMRI results in spatially mutually independent regions of the brain which are involved in a specific function. Since the true number of components is unknown, choosing the number of components allows a trade-off between size and number of ROIs. For this study, we used timer series which were already extracted and are publicly available through HCP consortium. For extracting this time series, between Group-PCA of 820 subjects were generated by MELODIC's incremental Group-PCA and then fed into group-ICA using FSL's MELODIC tool for dimensionality of 100 (Smith et al., 2013). It is worth noting that not all ICA components are necessarily physiologically meaningful, some of them may have been caused by head movement or cardiac process, but considering that we the 'bad' components were already extracted from the data by FIX-ICA (Salimi-Khorshidi et al., 2014), all the 200 components are expected to be carrying a large extent physiological information. ICA-200 covers cortical, sub cortical and cerebellum regions of the human brain. As the RSN assignment of nodes are unknown, we discard local examination of changes for this specific parcellation scheme. This parcellation scheme was formed using grayordinates available via HCP consortium and hence is considered as a surface-based parcellation.

- **Power** is a data-driven atlas which is formed by same-sized spheres around the putative functional coordinates (Power et al., 2011). These putative coordinates are combination of task-related and resting-state regions of the human brain. Through meta-analysis of more than 72 studies, 152 functional putative coordinates related to task fMRI were found. Further, 193 coordinates were also found by analysing a healthy resting-state cohort of 1000 subjects. By removing the overlaps between the coordinates found by these two methods, 264 coordinates remains as representatives of the functional activities within cortical, sub cortical and cerebellum of the human brain. This brain parcellation has a relatively small and same-sized regions of interests which helps avoid the possible overlaps between signals of different regions. Regions of interest in Power2011 is divided into 12 RSN networks including Sensory, Cingulo-opercular task control, Auditor, Default Mode Network, Ventral Attention Network, Visual, Fronto-parietal task control, Salience network, subcortical network, Cerebellar network and Dorsal Attention Network.

- **Gordon** is data-driven surface-based atlas which is comprised of 165 ROIs per hemisphere. The atlas was formed by boundary-mapping technique using voxel-wise seed-based correlations. The atlas is comprised of 12 distinct resting-state communities including Auditory, Cingulo-Opercular (CinguloOperc), Cingulo-Parietal, Default, Dorsal Attention (DorsalAttn), Fronto-Parietal, RetrosplenialTemporal, Senory Motor - Hand (SMhand), Sensory Motor - Mouth (SMmouth), Salience, Ventral Attention (VentralAttn) and Visual. Also, total of 47 parcels were grouped as 'None' meaning that the belong to none of the resting-state communities. The author presented a thorough validation and comparison analysis of the atlas with several other atlases, including the ones in this work, and conclude that Gordon's atlas is the most homogeneous in term of the parcel size and shapes (Gordon et al., 2016; Cohen et al., 2008). This atlas was recommended to be used on subject level although, for consistency, we use its group-level parcellation scheme.

- **Multimodal Parcellation (MMP)** is a surface-based parcellation scheme based on a combination of task fMRI, resting fMRI, Diffusion Tensor Imaging and structural MRI using 210 healthy subjects (Glasser et al., 2016). In MMP, each hemisphere is parcellated into 180 non-overlapping areas. We only use this parcellation scheme to confirm that the association between the ROI size and autocorrelation is also present in surface-based analysis. For further analysis, we will not present of MMP parcellation scheme.

#### S4.2 Resting-State Functional Connectivity

Using atlases discussed in section S4.1, we parcellate scan of each subject into set of ROIs. The temporal association between each pair ROIs is estimated via standardised Pearson's correlation coefficients, $\hat{Z}_{XY}$. For Naive Z-scores We use Eq. S5 and normalise the Fisher's $\hat{F}_{XY}$ with $1/\sqrt{N-3}$, where $N$ is the nominal degrees of freedom, to obtain the standardised statistics (i.e. $Z$-score). Similarly, for BH-corrected Z-scores, we use $1/\sqrt{\hat{N}}$, where $\hat{N}$ is the effective degrees of freedom and is calculated using Eq 9 of the main text. Finally, we obtain xDF-corrected Z-scores via Eq 11.

We then compare the Z-scores of different methods (shown in Figure 6 Figure 7 and Figure 8 of the main text and Figure S7-S17 of this document) to investigate how the choice of estimators may effect the interpretation of results for both statistical and density thresholding methods.

#### S4.3 Statistical Thresholding

In statistical thresholding (ST) we test each correlation coefficient against the null hypothesis that the observed correlation coefficient is equal to zero. This results in as many p-values as number of edges, $E = \frac{K(K-1)}{2}$, and therefore a correction for multiple testing should be applied. Among several of these correction methods, Bonferroni and FDR has been widely used in the field. The former uses a stringent threshold of $\alpha$-level/$E$ and controls the familywise error, the chance of one or more false positives; the latter uses uses the distribution of p-values to estimate an adaptive threshold, which controls the false positives as a proportion of the number of significant tests. Bonferroni can be severely conservative and FDR somewhat conservative in the presence of dependence between the $E$ tests (Benjamini et al., 2006), but FDR has been commonly used in network studies, such as Geerligs et al. (2014) and Ridley et al. (2015).Finally, any correlation coefficients survived the hypothesis test and the multiple comparison error correction, is set to one and any of them that failed to reject the null is set to zero. However note that FDR is adaptive and subject-dependent thresholds may be troublesome for further network analysis.

#### S4.4 Proportional Thresholding

Proportional thresholding (PT) is a widely used method for thresholding adjacency matrices over a range of densities where densities are defined as number of edges over all possible number of edges. In DT, edge weights of an adjacency matrix is thresholded by a certain value, such that supra-threshold edges are preserved while the rest is set to zero. In this setting, only strongly connected edges are preserved. It is worth noting that one should be careful with selecting the boundaries as setting the threshold too high may results in too many leaves (i.e. isolated nodes) while a low threshold may lead to a lattice. Alternatively, functional connectivities may be thresholded on Cost-Efficient (CE) density, where the network demonstrate the highest global efficiency while preserving the lowest wiring cost (i.e. density). In contrast to statistical thresholding, the proportional thresholding enjoys an universal density across all subjects. For the reminder of this work, when we refer to PT it means the network is thresholded proportionally on the CE density.

#### S4.5 Graph Theoretical Measures

To demonstrate that how accounting for autocorrelation may effect the interpretation of topological features, we use weighted degrees and betweenness as measures of centrality, local and global efficiencies as measures of segregation and finally, their combination as measure of complexity.

##### S4.5.1 Degree and Betweenness Centrality

Degree and betweenness centrality might be of simplest graph theoretical measures to summaries the centrality measures of a node. In degree centrality, we accumulate all the weights connected to a node as strength or weighted degree of a node (Rubinov and Sporns, 2010a). In Betweenness centrality, we measure how often a node appears in a shortest path between all pair of nodes (Brandes, 2001). In the next section, we explain the concept of shortest path lengths.

##### S4.5.2 Measure of Integration

Integration in a network can be mathematically quantified using the shortest path between two nodes in a topological description of the connectome. In graph theory terminology, a *Path* is a collection of nodes and edges which represent the shortest topological distance between two specific nodes (Rubinov and Sporns, 2010a). The average shortest Path between all nodes of a network is called the *characteristic path length* ($L$) (Watts and Strogatz, 1998). A smaller characteristic path length, suggests a higher level of integration within a network. Throughout this work we use Path Length for Characteristic Path Length, unless otherwise stated. The characteristic path length can be quantified as:

$$L = \frac{1}{V} \sum_{i \in V} \frac{\sum_{j \in N, j \neq i} \sum_{a_{ln} \in g_{i \leftrightarrow j}} a_{uv}}{V - 1} \tag{S12}$$

where $N$ is number of nodes, $a_{uv}$ is the numnber of the edge between node $u$ and $v$ in case of binary local efficiency. Note that one can calculate weighted local efficiency by replacing $a_{uv}$ with $w_{uv}$ which is the weights of edges between the two nodes $u$ and $v$. According to Eq. S12 if a node happens to be isolated, its shortest path turns out to be infinity which leads Eq. S12 to be infinity too. To address this issue, Latora and Marchiori (2001) proposed a similar method, called global efficiency, which is an average of the inverse shortest path, therefore, the shortest path of isolated nodes is zero. Eq. S12

##### S4.5.3 Measures of Segregation

The level of segregation in a network can be measured as to what extent a nodes of a network can be clustered or grouped. To quantify this notion, Latora and Marchiori (2001) proposed a measure of local segregation, called local efficiency which, for weighted networks, can be formulated as (Rubinov and Sporns, 2010b):

$$E_i = \frac{\sum_{i,h \in V, j \neq i} \left( w_{ij} w_{ih} (d_{jh}(V_i))^{-1} \right)^{1/3}}{k_i(k_i - 1)} \tag{S13}$$

where $d_{jh}(v_i)$ is number of edges of the shortest path between nodes $j$ and $h$ that are only one edge away from node $i$. For binary networks, the local efficiency is defined as,

$$E_i = \frac{\sum_{i,h \in V, j \neq i} a_{ij} a_{ih} (d_{jh}(V_i))^{-1}}{k_i(k_i - 1)} \tag{S14}$$

Local efficiency is tightly similar to clustering coefficients, but they are not identical as the latter measures the probability of formation of a triangle around node $i$, while the former measures the topological distance between two neighbours of node $i$.

### S5 Results

#### S5.1 Bias of Standard Error for the Fisher Transformation

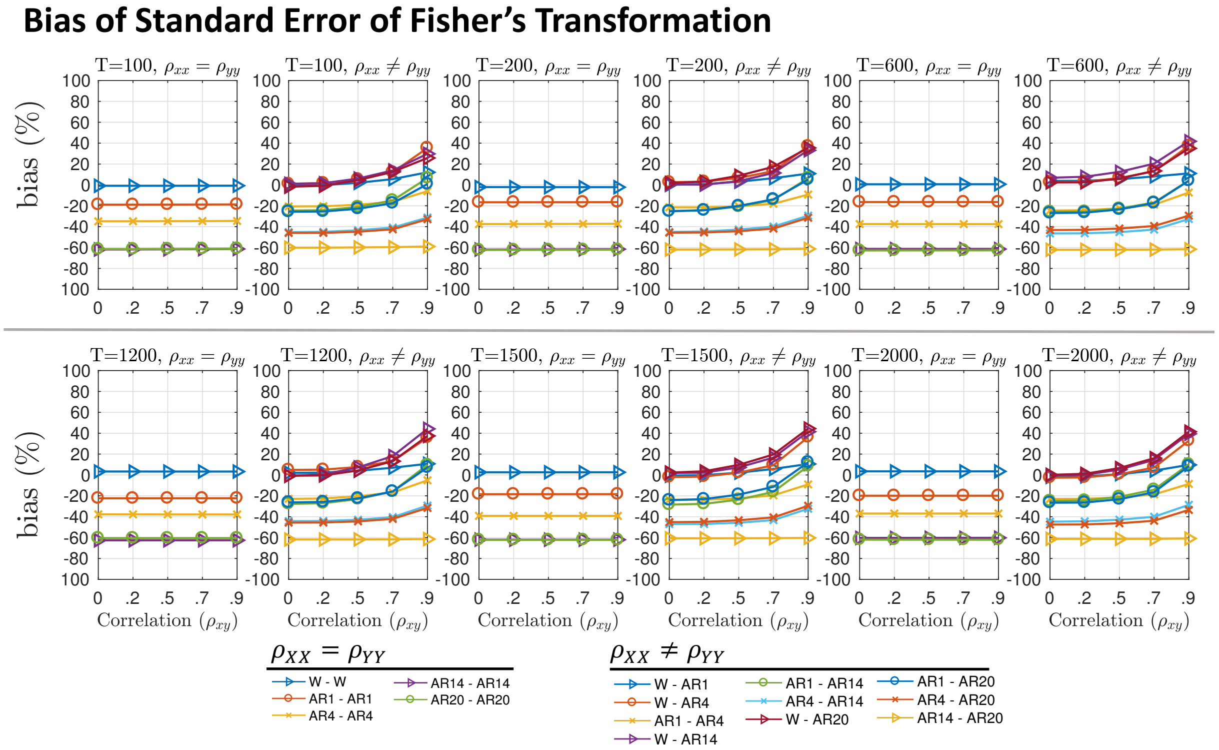


Figure S2: Biases in standard error of Fisher's transformed sample correlation coefficients ($F_{XY}$) obtained by comparing the theoretical standard error $\sigma_{F_{XY}} = \frac{1}{\sqrt{N-3}}$ against the standard error of Monte-Carlo simulated correlations. It is important to note that the we use the nominal degrees of freedom in these simulation, $N$, to reflect on the severe effect of autocorrelation in standard error as well as failure of the Fisher's transformation to stablise the standard error.

#### S5.2 Oracle Simulation

Figure S3.A shows the biases of oracle simulations for xDF, for different sample sizes and autocorrelation structures (similar to Figure 4 of the main text). The result suggest that the bias xDF never exceed the [-10 10] bias range, with higher biases on short time series (T=100) and almost zero biases on longer time series (T=2000). Similar analysis for HB suggests that, even on Oracle simulations, the statistical aliasing effect biases the estimator once the correlation coefficient is increased. Similar to the Monte Carlo simulations (Figure 4.B & Figure 4.C), this effect is invariant of the sample sizes.

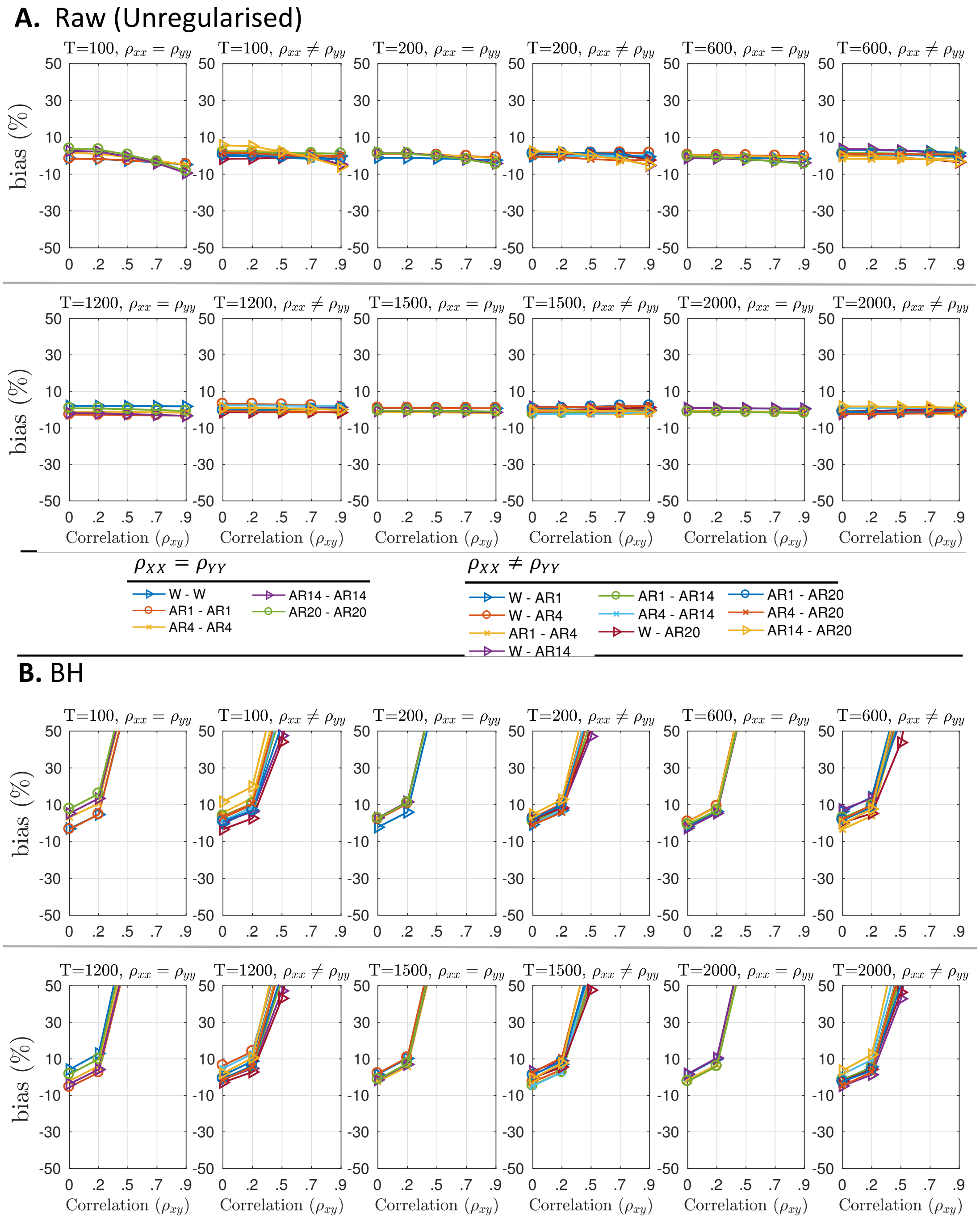


Figure S3: Biases of Oracle simulations. These simulation were performed following Algorithm 1 and percentage biases were calculated via Eq. S10. **Panel A,** illustrates the biases in standard deviation of the xDF estimates across 6 sample sizes without any form of tapering. **Panel B,** similar results for BH correction method, without any form of regularisation on the ACF. Note that we do not use any form of regularisations for methods investigated here since the true parameters are used and therefore the calculations are noise-free.

##### S5.3 False Positive Rates

Figure S4 illustrates the FPR of each method for a given $\alpha-$level, complimenting Figure 3.C of the main text for 1% and 5% $\alpha-$level.

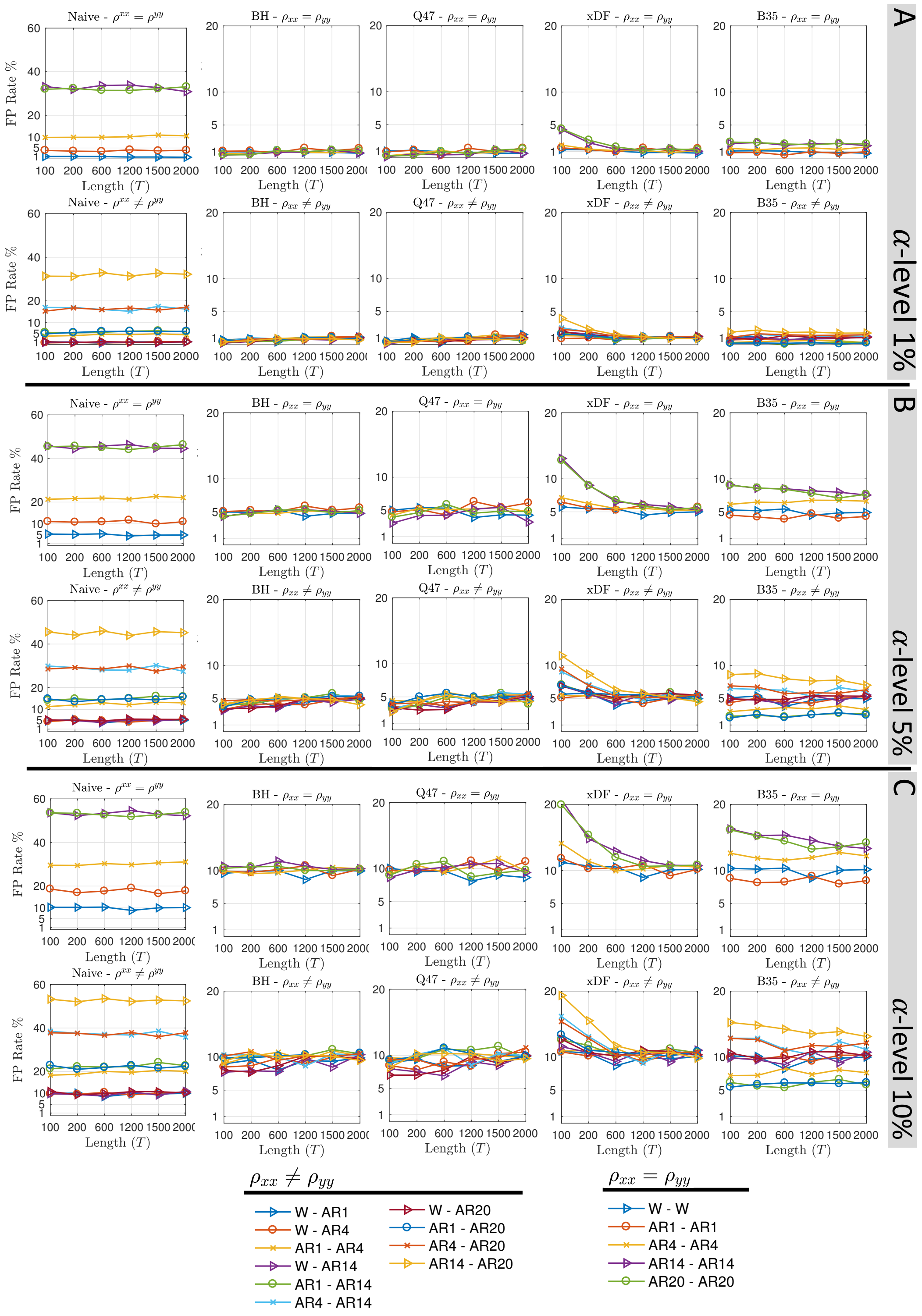


Figure S4: False positive rate analysis of the pair-wise Monte Carlo simulations. In each panel, the FPR of each method (i.e. columns) for identical and different AC structures (i.e. rows) were illustrated. **Panel A,** False positives of $\alpha-$level=1%. **Panel B,** False positives of $\alpha-$level=5%; this is identical to Figure 3.C of the main text. **Panel C,** False positives of $\alpha-$level=10%.

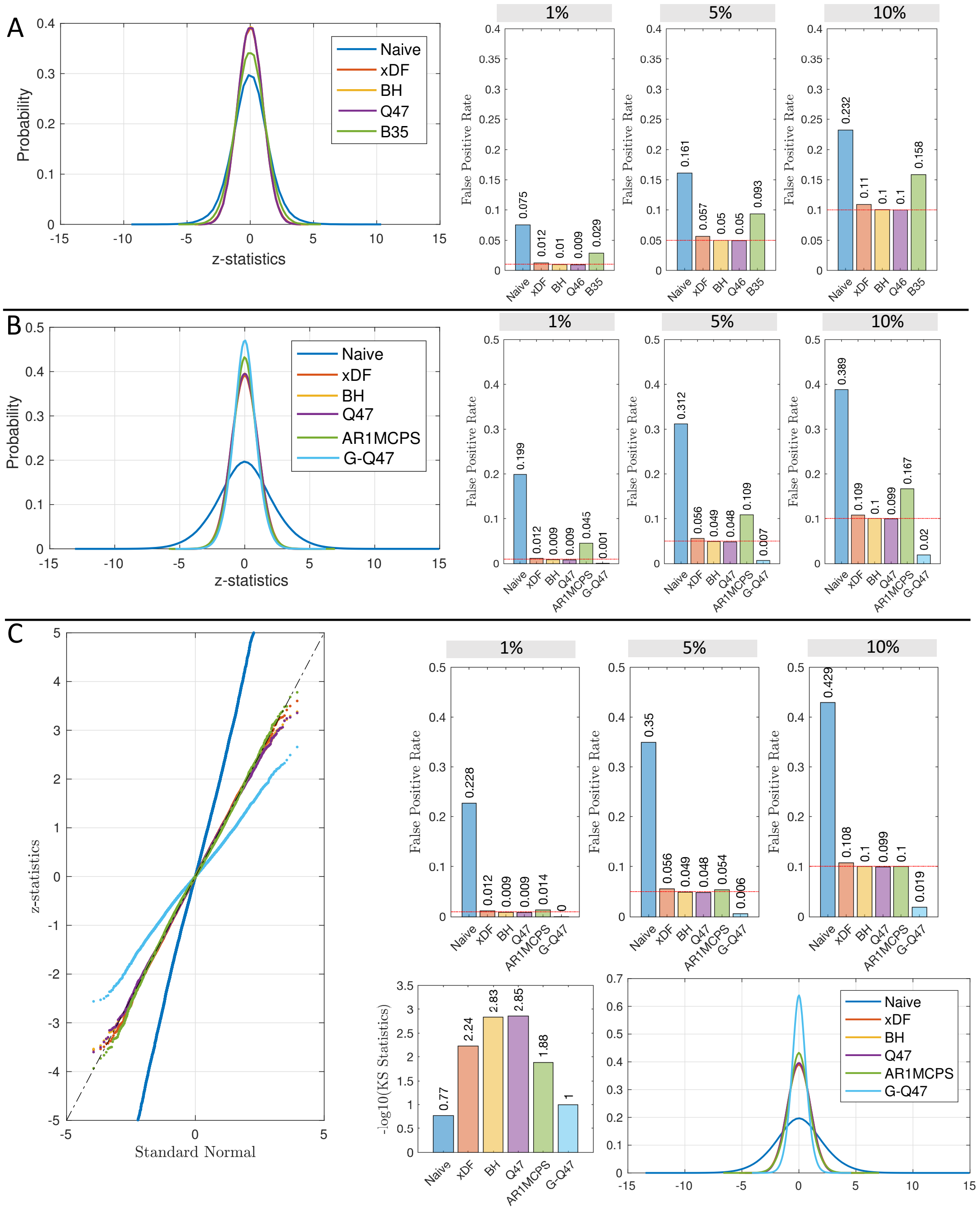


Figure S5: **Panel A,** illustrates the pair-wise inter-subject scrambling across five correction methods. The histograms of the z-statistics of ISS are shown on left and the FPRs of $\alpha-$levels$=\{1\%, 5\%, 10\%\}$ are shown on right. **Panel B,** FPR and KS analysis of the simulated correlation matrices (AC structure from 118528) for six correction methods; including the global methods (G-Q47 and AR1MCPS). The panel follows a similar layout as panel A. **C.** illustrates the same analysis but for AC structures inherited from subject 135932. For more information regarding the simulated correlation matrices, see Section S3.5

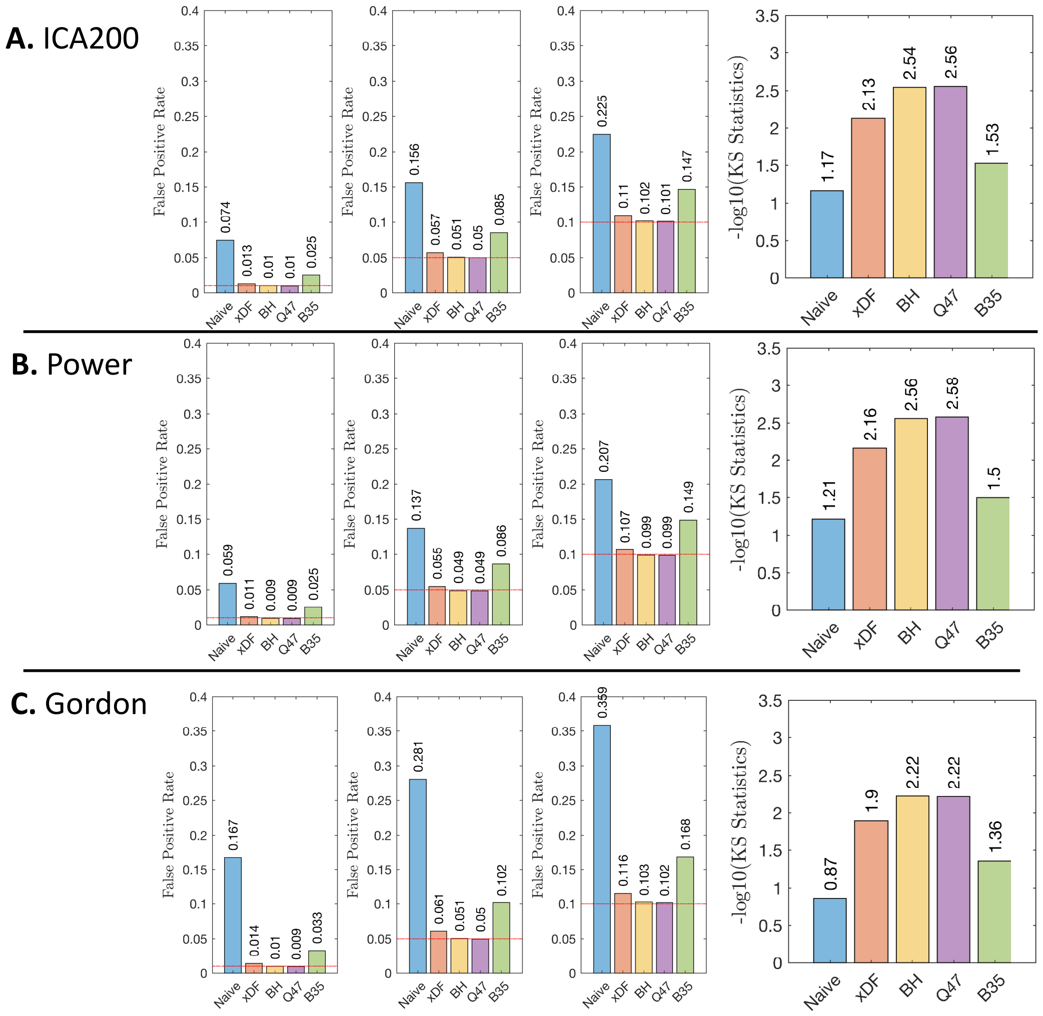


Figure S6: Inter-subject scrambling for **A** ICA200, **B** Power atlas and **C** Gordon atlas. Figure 3.A in the main text illustrators similar results for Yeo Atlas.

#### S5.4 Specificity, Sensitivity and ROC Curves

--

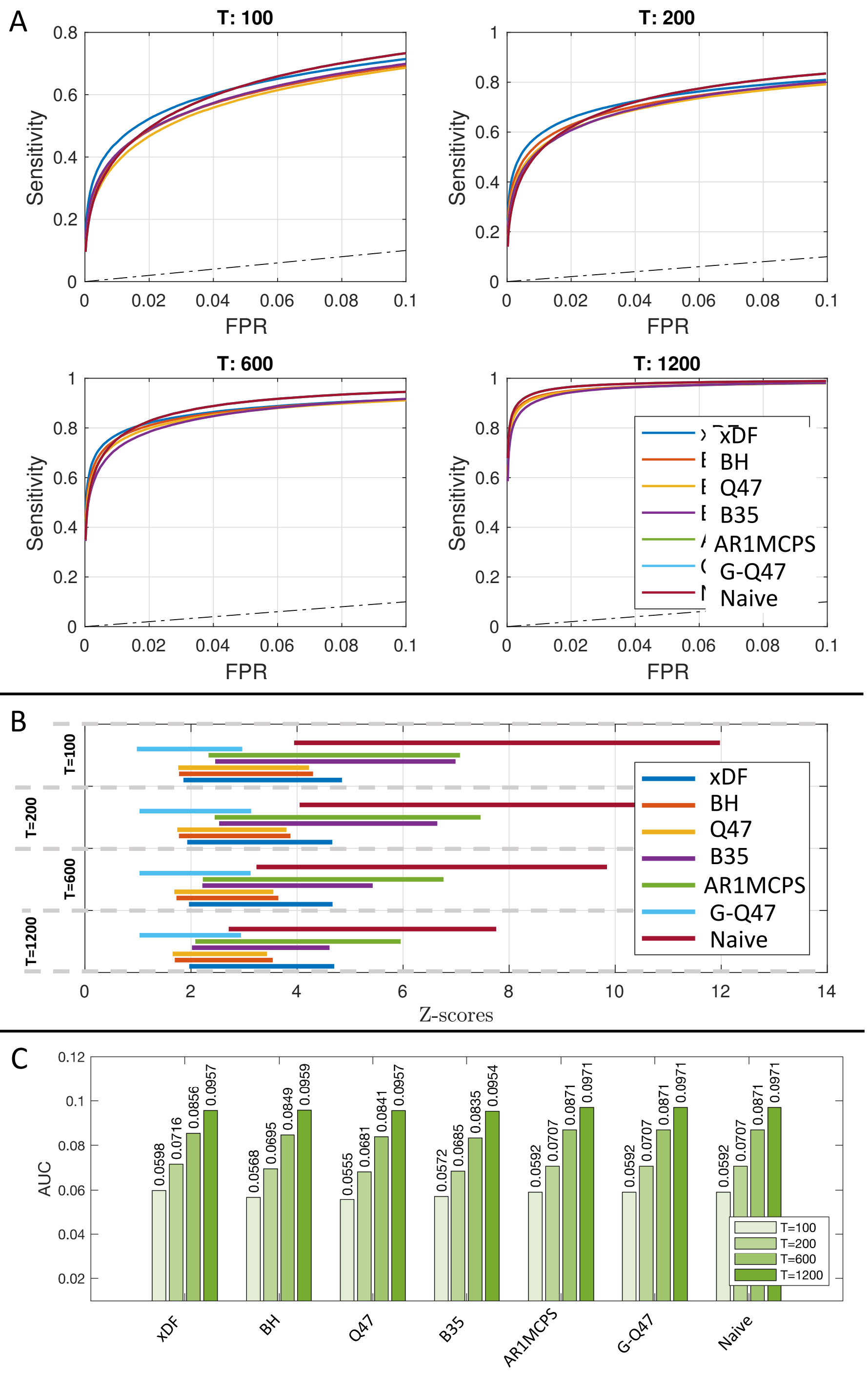


Figure S7: **Panel A,** shows the ROC curves for each of the seven methods across different sample sizes; $T100, 200, 600, 1200$. The reference line was shown as a dashed black line. **Panel B,** shows the threshold ranges required for each method to obtain 0 to 10% FPR. **Panel C,** Shows the Area Under the Curve (AUC) for the ROC curves illustrated in panel A.

#### S5.5 Monte Carlo Simulations for Different Tapering Methods

--

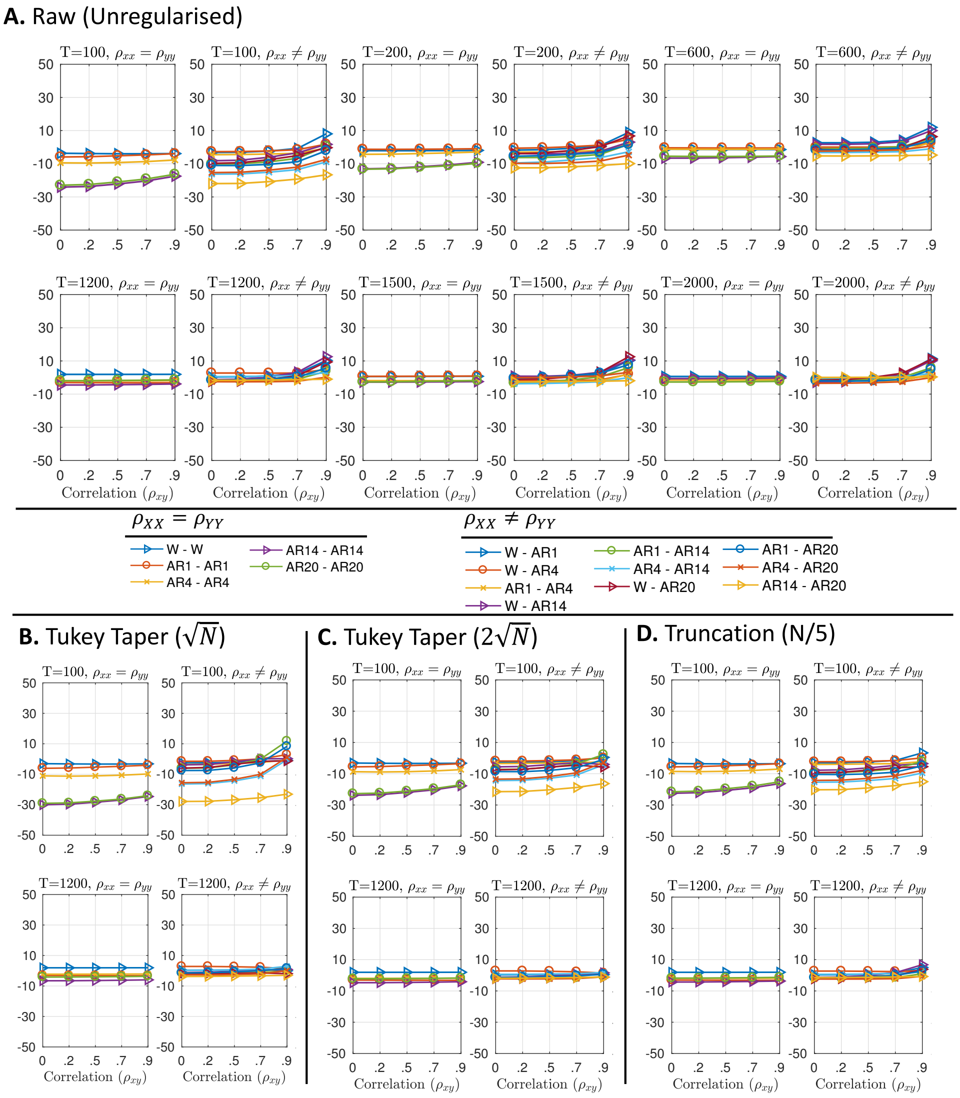


Figure S8: The bias of standard deviation for xDF with different tapering techniques. **Panel A,** without any form of tapering **Panel B,** with Tukey tapering $M = \sqrt{N}$. **Panel C,** with Tukey tapering $M = 2\sqrt{N}$. **Panel D,** with truncation $M = N/5$. For details of the simulation see Algorithm 2, and Eq. S10 for the bias calculation.

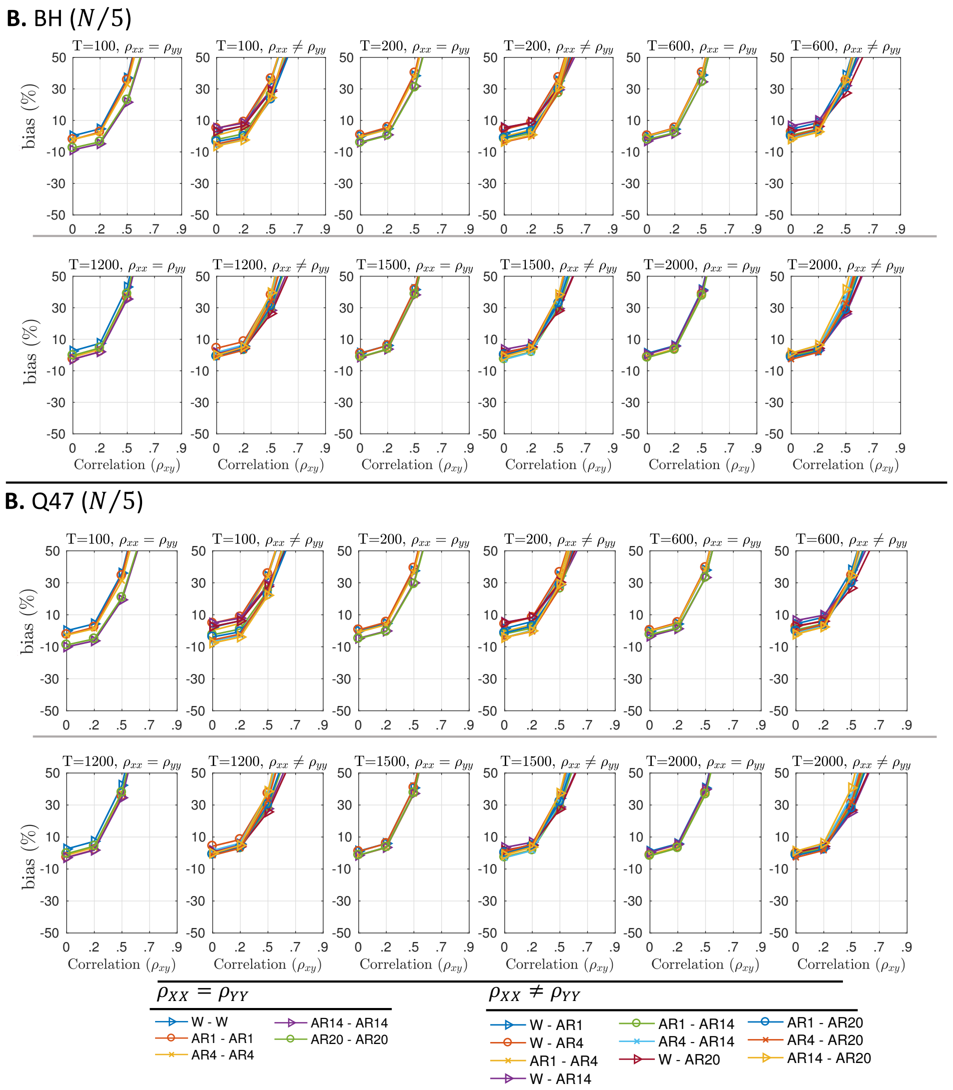


Figure S9: The biases in standard deviation estimates in BH and Q47 with $\frac{N}{5}$ truncation factor. **Panel A,** BH with $M = \frac{N}{5}$ truncation . **Panel B,** Q47 with $M = \frac{N}{5}$ truncation. For details of the simulation see Algorithm 2, and Eq. S10 for the bias calculation.

##### S5.6 Changes in Functional Connectivity

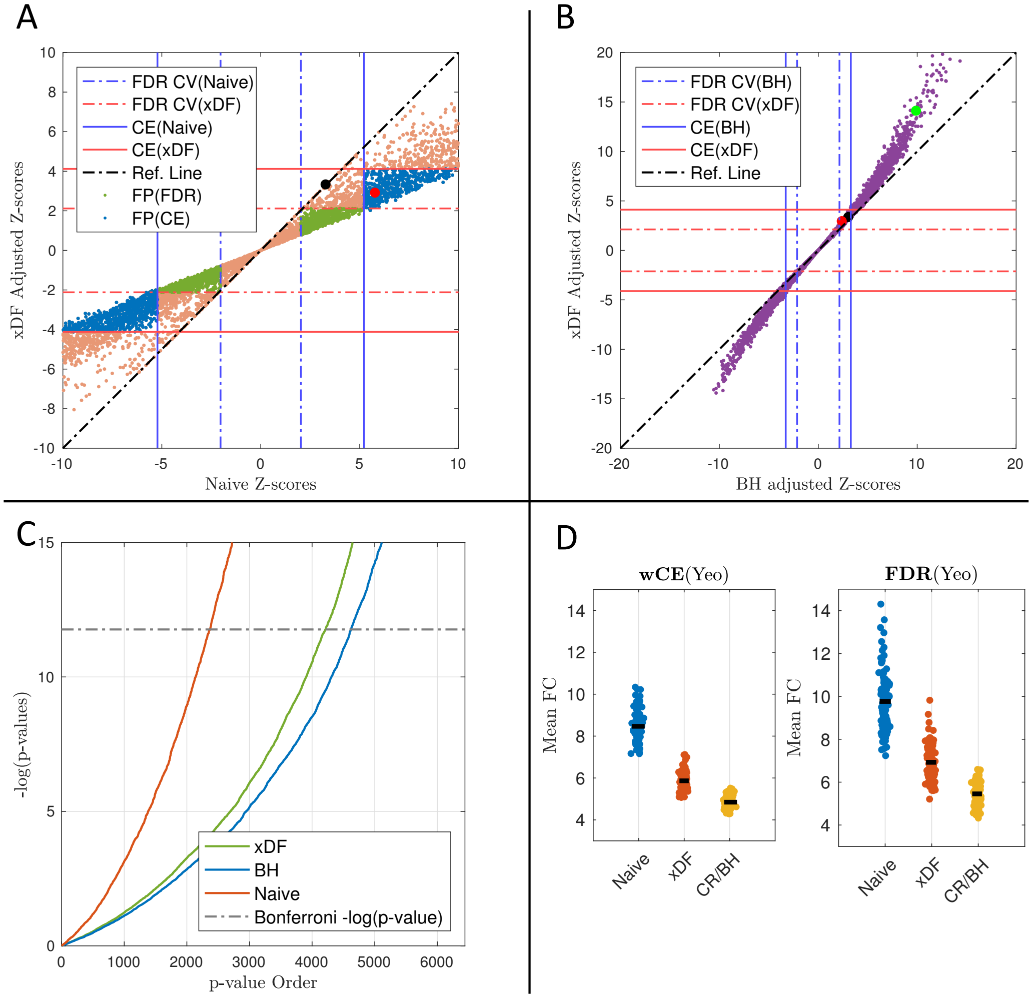


Figure S10: Changes in rsFC of HCP 135932 due to Naive, xDF and BH corrections. The figure layout is identical to Figure 6 of the main text. Panel **A,** illustrates the changes in FC strengths of xDF corrected connectivity maps, relative to the Naive correction. Solid line shows the critical value corresponding to the cost-efficient density. Dashed lines illustrates the critical values of FDR-corrected p-values. The false positive edges were colour coded as green (FPR in CE) and blue (FPR in FDR). the black dot indicate an edge between nodes 37 and 87 and the red dot indicate the edge between nodes 7 and 85. Panel **B,** shows the changes in FC strengths with xDF and BH corrections. The green dot indicates the edge between nodes 88 and 23. The remaining are similar to the settings in Panel A. Panel **C,** shows the changes in p-values of Naive correction relative to xDF (orange dots) and xDF relative to BH (purple dots). Panel **D,** shows the overall changes mFC of for each correction method.

#### S5.7 Changes in Graph Theoretical Measures due to Autocorrelation Correction

##### S5.7.1 Un-thresholded Networks

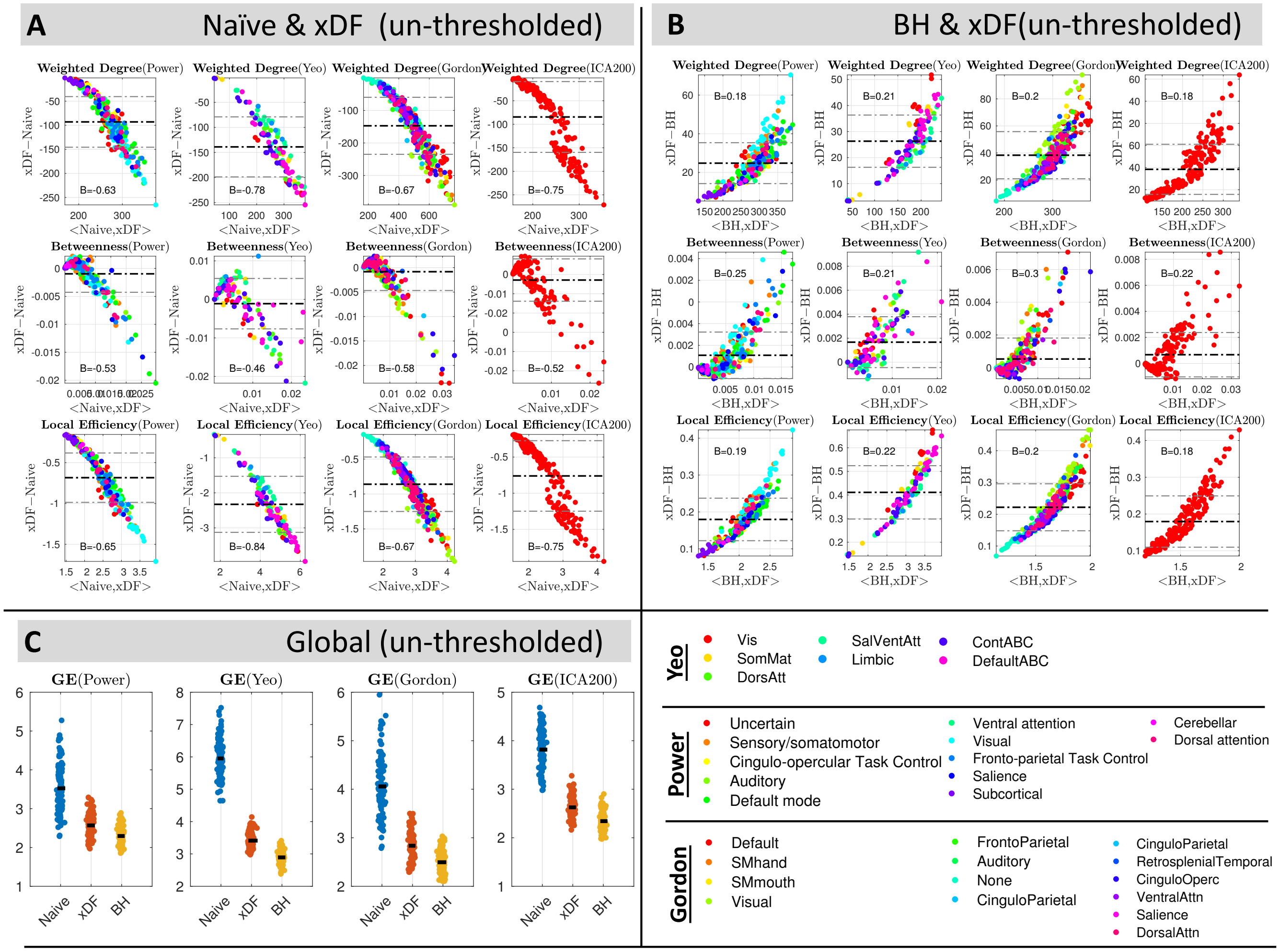


Figure S11: Changes in graph theoretical measures of unthresholded functional connectomes. **Panel A,** illustrates the changes after correcting the functional connectivities via xDF in weighted degree, betweenness centrality and local efficiency (i.e. rows) across three atlases (columns; Power, Yeo and Gordon). **Panel B,** illustrates the changes dues to aliasing effect with an identical layout to panel A. **Panel C,** shows the change in global efficiency of the subjects parcelled with Power (left), Yeo (middle) and Gordon (right) for three correction methods.

##### S5.7.2 Proportional- and Statistical- Thresholded Weighted Networks of Gordon, Power and ICA200 Atlas

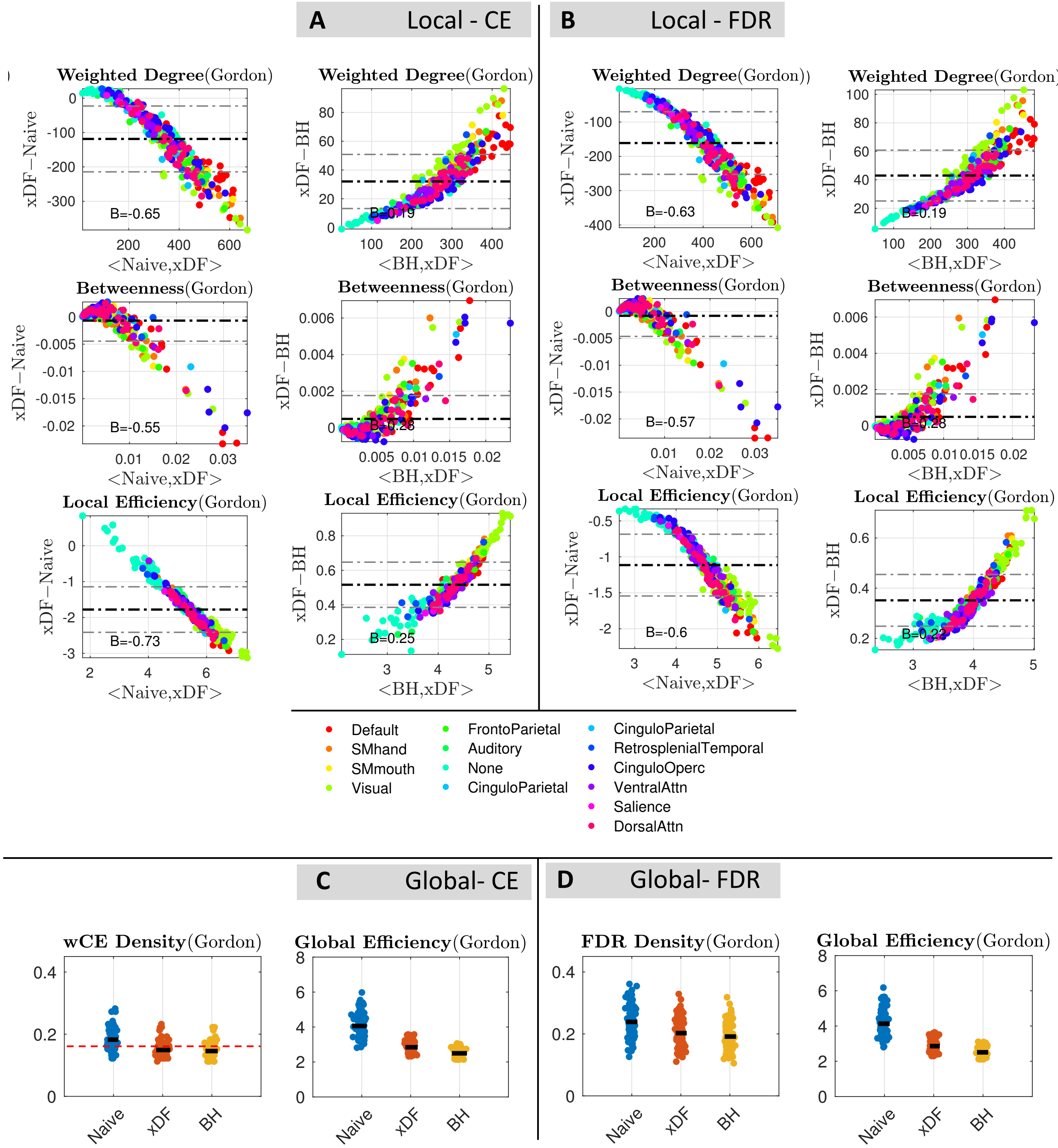


Figure S12: Overall changes in global and local graph theoretical measures of HCP unrelated subjects parcellated with the Gordon atlas. The layout in this Figure is identical to of Figure 7. **Panel A,** Bland Altman plots of the changes in weighted (top), betweenness (middle) and local efficiency (bottom) of 99 subjects thresholded proportionally on CE density. The nodes are colour coded according to their resting-state network assignment. **Panel B** Shows the similar graph measures, but with statistical thresholding (corrected via FDR correction). **Panel C** shows the changes in weighted CE density (left) and Global efficiency (right), similarly, **Panel D** illustrates the same results for networks thresholded via FDr-based statistical thresholding

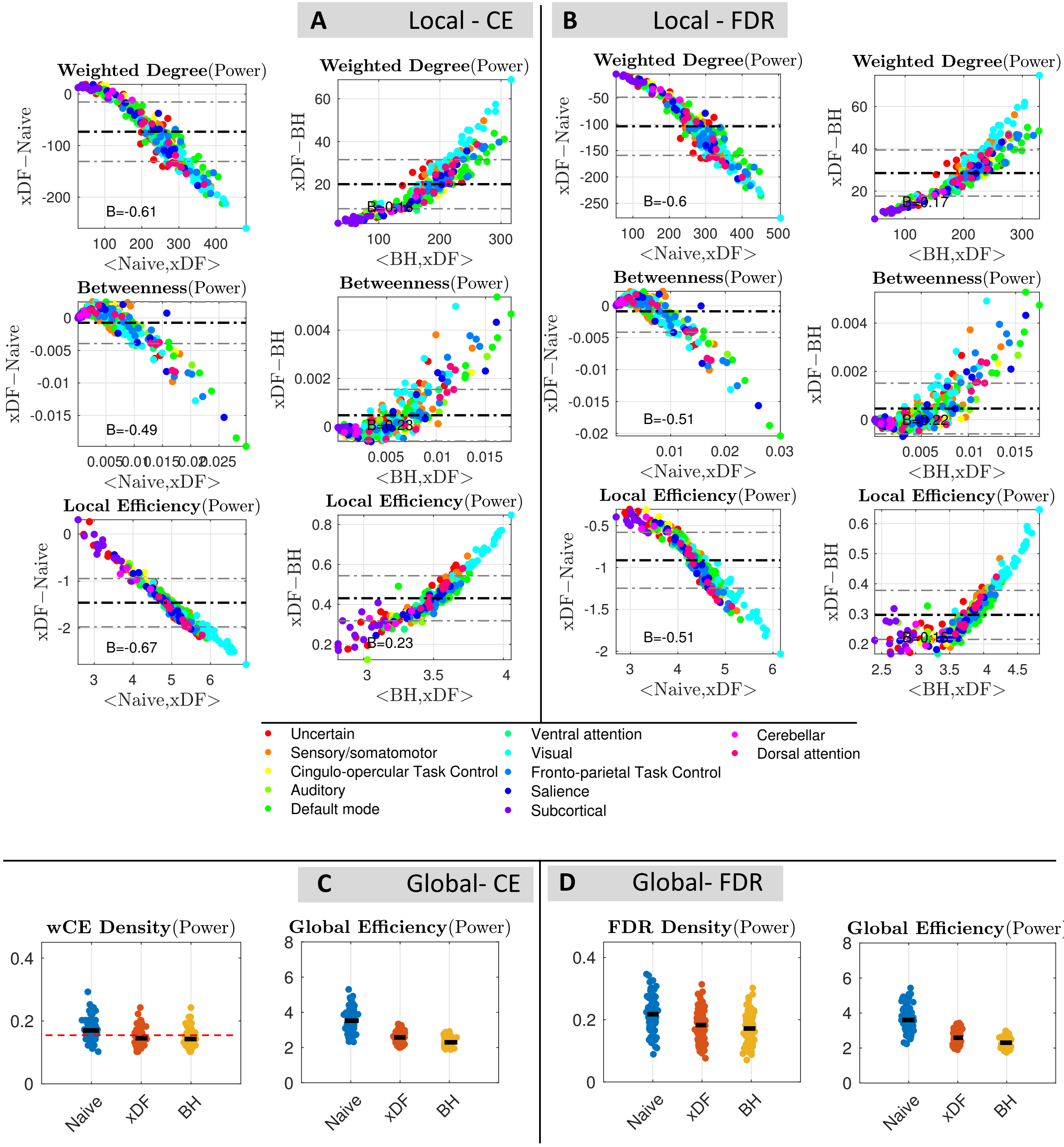


Figure S13: Overall changes in global and local graph theoretical measures of HCP unrelated subjects parcellated with the Power atlas. The layout in this Figure is identical to of Figure 7. **Panel A,** Bland Altman plots of the changes in weighted (top), betweenness (middle) and local efficiency (bottom) of 99 subjects thresholded proportionally on CE density. The nodes are colour coded according to their resting-state network assignment. **Panel B** Shows the similar graph measures, but with statistical thresholding (corrected via FDR correction). **Panel C** shows the changes in weighted CE density (left) and Global efficiency (right), similarly, **Panel D** illustrates the same results for networks thresholded via statistical thresholding and corrected with FDR.

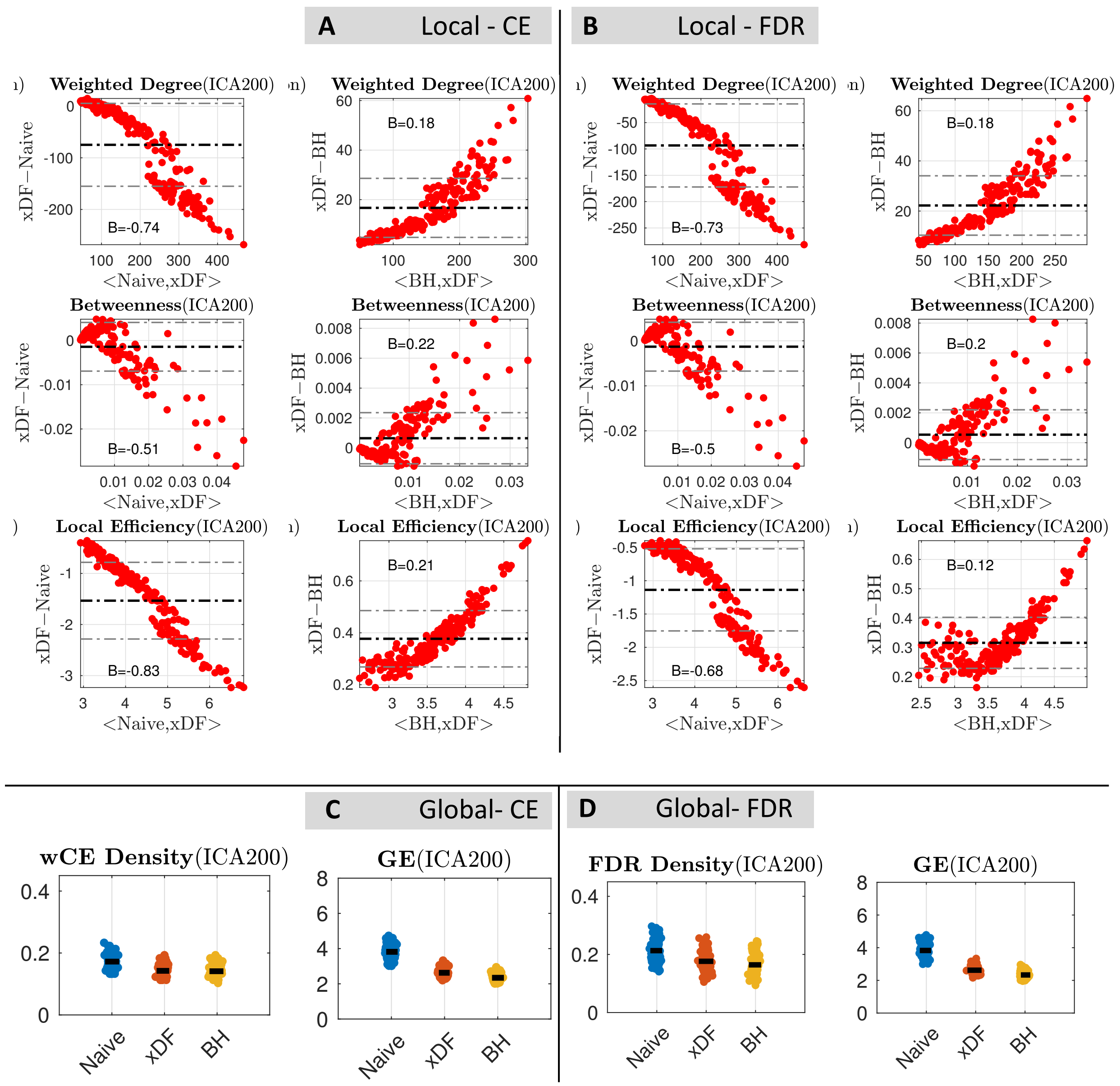


Figure S14: Overall changes in global and local graph theoretical measures of HCP subjects parcellated by ICA200. The layout in this Figure is identical to of Figure 7. **Panel A,** Bland Altman plots of the changes in weighted (top), betweenness (middle) and local efficiency (bottom) of 99 subjects thresholded proportionally on CE density. The nodes are colour coded according to their resting-state network assignment. **Panel B** Shows the similar graph measures, but with statistical thresholding (corrected via FDR correction). **Panel C** shows the changes in weighted CE density (left) and Global efficiency (right), similarly, **Panel D,** illustrates the same results for networks thresholded via statistical thresholding and corrected with FDR.

Table S2: Proportion of nodes, in Power atlas, affected in Naive vs. xDF correction of weighted rsFC

| Thresholding | Weighted Degree | Weighted Betweenness | Local Efficiency |
|---|---|---|---|
| PT (CE) | 71.97% | 15.53% | 93.20% |
| ST (FDR) | 83.33% | 17.80% | 98.11% |

Table S3: Proportion of nodes, in Gordon atlas, affected in Naive vs. xDF correction of weighted rsFC

| Thresholding | Weighted Degree | Weighted Betweenness | Local Efficiency |
|---|---|---|---|
| PT (CE) | 73.87% | 2.10% | 94.30% |
| ST (FDR) | 87.69% | 36.03% | 98.50% |

Table S4: Proportion of nodes, in Power atlas, affected in xDF vs. BH correction of weighted rsFC

| Thresholding | Weighted Degree | Weighted Betweenness | Local Efficiency |
|---|---|---|---|
| PT (CE) | 5.30% | 0% | 85.60% |
| ST (FDR) | 14.40% | 0% | 83.71% |

Table S5: Proportion of nodes, in Gordon atlas, affected in xDF vs. BH correction of weighted rsFC

| Thresholding | Weighted Degree | Weighted Betweenness | Local Efficiency |
|---|---|---|---|
| PT (CE) | 31.53% | 2.7% | 93.40% |
| ST (FDR) | 42.94% | 2.7% | 93.39% |

##### S5.7.3 Proportional- and Statistical- Thresholded Binary Networks of Gordon, Power and ICA200 Atlas

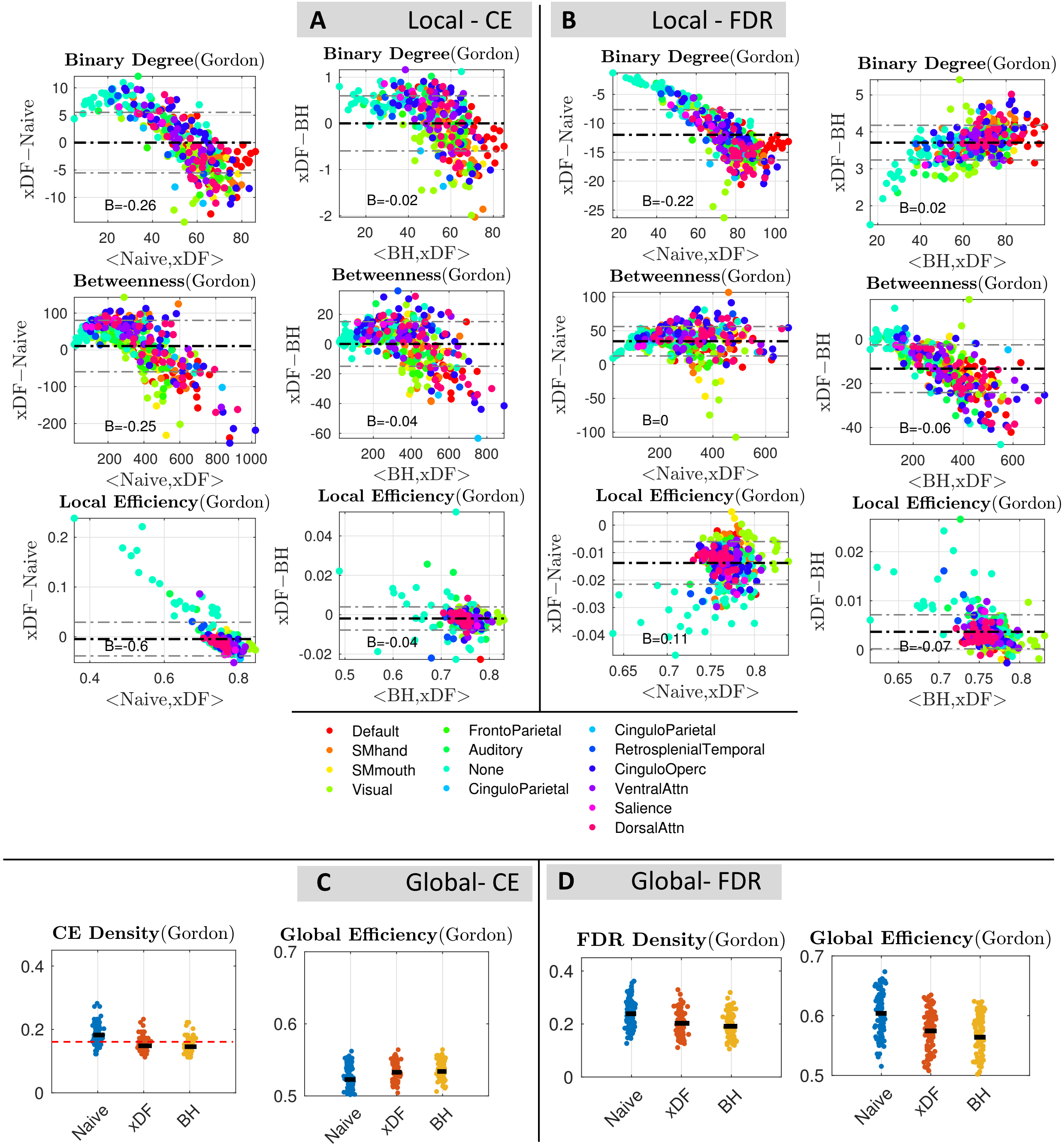


Figure S15: Overall changes in global and local graph theoretical measures of HCP unrelated subjects parcellated with the Gordon atlas. The layout in this Figure is identical to of Figure 7. **Panel A,** Bland Altman plots of the changes in binary (top), betweenness (middle) and local efficiency (bottom) of 99 subjects thresholded proportionally on CE density. The nodes are colour coded according to their resting-state network assignment. **Panel B** Shows the similar graph measures, but with statistical thresholding (corrected via FDR correction). **Panel C** shows the changes in weighted CE density (left) and Global efficiency (right), similarly, **Panel D** illustrates the same results for networks thresholded via FDr-based statistical thresholding

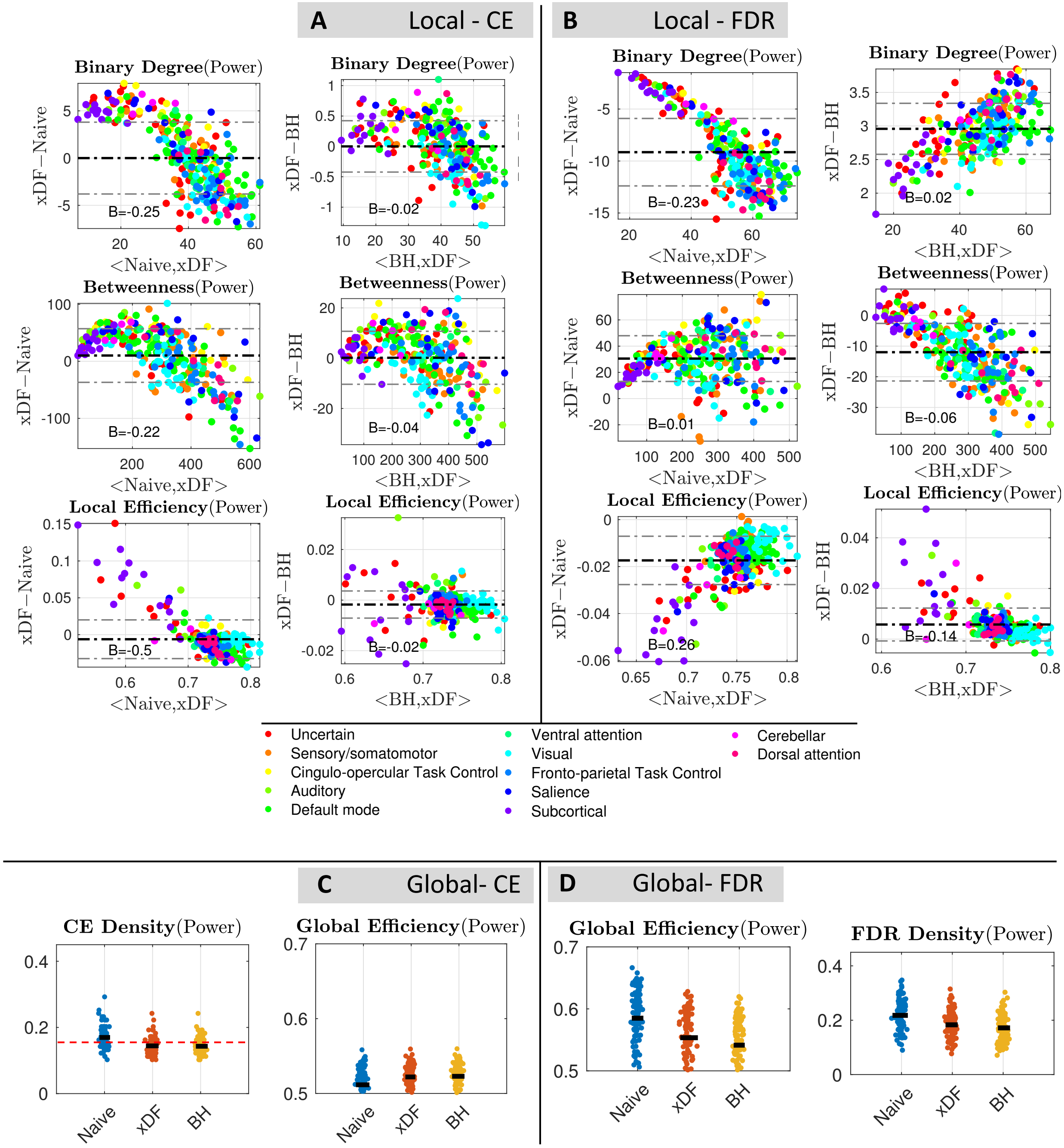


Figure S16: Overall changes in global and local graph theoretical measures of HCP unrelated subjects parcellated with the Power atlas. The layout in this Figure is identical to of Figure 7. **Panel A,** Bland Altman plots of the changes in binary (top), betweenness (middle) and local efficiency (bottom) of 99 subjects thresholded proportionally on CE density. The nodes are colour coded according to their resting-state network assignment. **Panel B** Shows the similar graph measures, but with statistical thresholding (corrected via FDR correction). **Panel C** shows the changes in weighted CE density (left) and Global efficiency (right), similarly, **Panel D** illustrates the same results for networks thresholded via statistical thresholding and corrected with FDR.

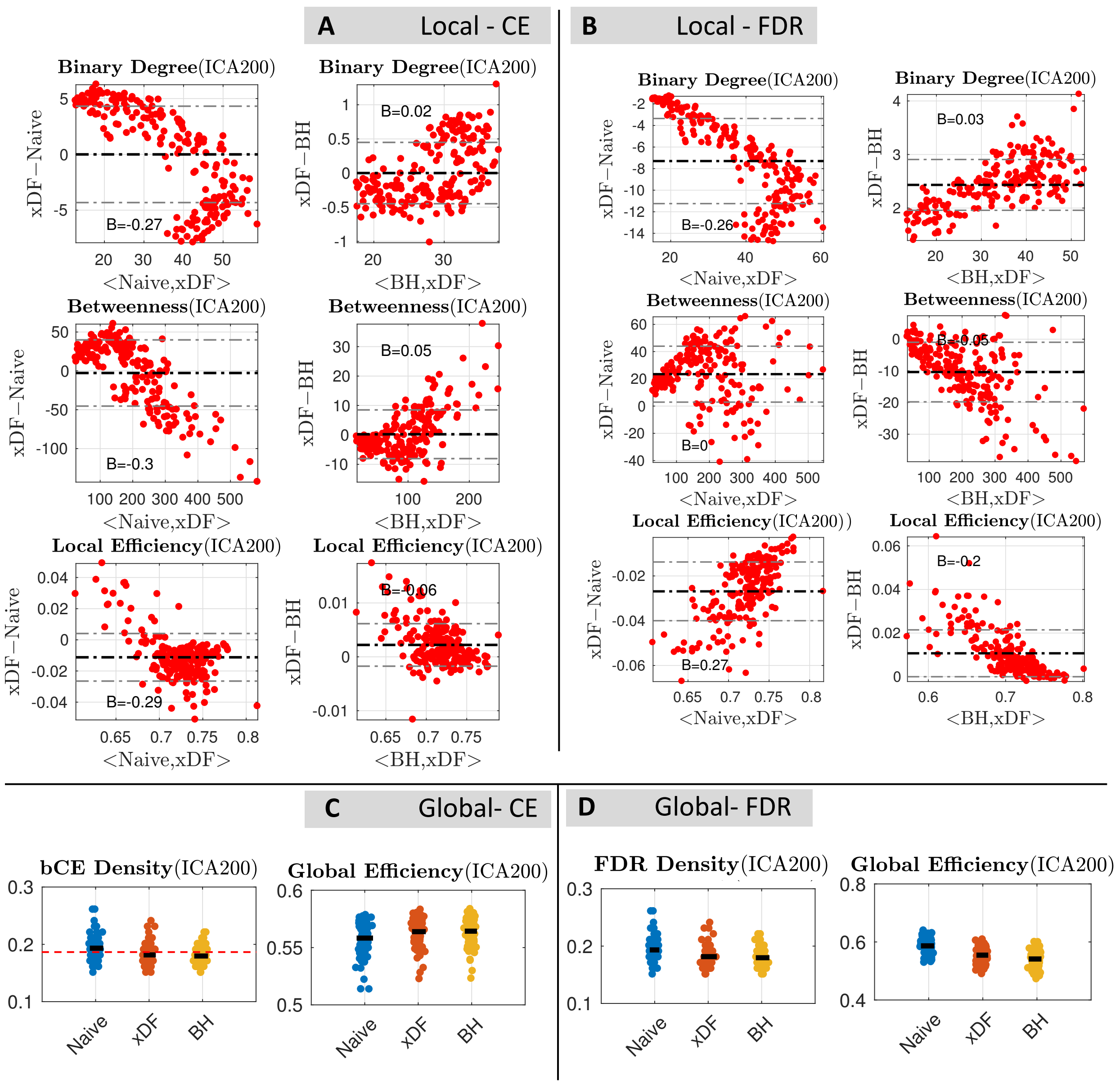


Figure S17: Overall changes in global and local graph theoretical measures of HCP subjects parcellated by ICA200. The layout in this Figure is identical to of Figure 7. **Panel A,** Bland Altman plots of the changes in binary (top), betweenness (middle) and local efficiency (bottom) of 99 subjects thresholded proportionally on CE density. The nodes are colour coded according to their resting-state network assignment. **Panel B** Shows the similar graph measures, but with statistical thresholding (corrected via FDR correction). **Panel C** shows the changes in weighted CE density (left) and Global efficiency (right), similarly, **Panel D,** illustrates the same results for networks thresholded via statistical thresholding and corrected with FDR.

Table S6: Proportion of nodes, in Power atlas, affected in Naive vs. xDF correction in binary rsFC

| Thresholding | Binary Degree | Binary Betweenness | Local Efficiency |
|---|---|---|---|
| PT (CE) | 71.96% | 15.53% | 93.18% |
| ST (FDR) | 73.10% | 0% | 0% |

Table S7: Proportion of nodes, in Gordon atlas, affected in Naive vs. xDF correction in binary rsFC

| Thresholding | Binary Degree | Binary Betweenness | Local Efficiency |
|---|---|---|---|
| PT (CE) | 73.87% | 33.03% | 94.29% |
| ST (FDR) | 87.08% | 0% | 0% |

Table S8: Proportion of nodes, in Power atlas, affected in xDF vs. BH correction in binary rsFC

| Thresholding | Binary Degree | Binary Betweenness | Local Efficiency |
|---|---|---|---|
| PT (CE) | 5.30% | 0% | 85.60% |
| ST (FDR) | 0% | 0% | 0% |

Table S9: Proportion of nodes, in Gordon atlas, affected in xDF vs. BH correction in binary rsFC

| Thresholding | Binary Degree | Binary Betweenness | Local Efficiency |
|---|---|---|---|
| PT (CE) | 31.53% | 2.70% | 93.39% |
| ST (FDR) | 0% | 0% | 0% |

### S6 Reproducibility

#### S6.1 Figures in the Main Text

Here we present sets of links which points out to codes we used to produce each figure in the paper;

- **Figure 1**: `https://github.com/asoroosh/xDF_Paper18/tree/master/Figure1/`
- **Figure 2**: `https://github.com/asoroosh/xDF_Paper18/tree/master/Figure2/`
- **Figure 3**: `https://github.com/asoroosh/xDF_Paper18/tree/master/Figure3/`
- **Figure 4**: `https://github.com/asoroosh/xDF_Paper18/tree/master/Figure4/`
- **Figure 5**: `https://github.com/asoroosh/xDF_Paper18/tree/master/Figure5/`
- **Figure 6**: `https://github.com/asoroosh/xDF_Paper18/tree/master/Figure6/`

- **Figure 7**: `https://github.com/asoroosh/xDF_Paper18/tree/master/Figure7/`
- **Figure 8**: `https://github.com/asoroosh/xDF_Paper18/tree/master/Figure8/`

##### S6.2 Figures in the Supplementary Material

- **Figure S1**: `https://github.com/asoroosh/xDF_Paper18/tree/master/SupplementaryMaterials/FigureS1`
- **Figure 2**: `https://github.com/asoroosh/xDF_Paper18/tree/master/SupplementaryMaterials/FigureS2`

The remaining figures were obtained using scripts used in Section S6.1.
